# The Energetics and Ion Coupling of Cholesterol Transport Through Patched1

**DOI:** 10.1101/2023.02.14.528445

**Authors:** T. Bertie Ansell, Robin A. Corey, Lucrezia Vittoria Viti, Maia Kinnebrew, Rajat Rohatgi, Christian Siebold, Mark S. P. Sansom

## Abstract

Patched1 (PTCH1) is the principal tumour suppressor protein of the mammalian Hedgehog (HH) signalling pathway, implicated in embryogenesis and tissue homeostasis. PTCH1 inhibits the Class F G protein-coupled receptor Smoothened (SMO) via a debated mechanism involving modulating accessible cholesterol levels within ciliary membranes. Using extensive molecular dynamics (MD) simulations and free energy calculations to evaluate cholesterol transport through PTCH1, we find an energetic barrier of ~15-20 kJ mol^-1^ for cholesterol export. In simulations we identify cation binding sites within the PTCH1 transmembrane domain (TMD) which may provide the energetic impetus for cholesterol transport. *In silico* data are coupled to *in vivo* biochemical assays of PTCH1 mutants to probe coupling between transmembrane motions and PTCH1 activity. Using complementary simulations of Dispatched1 (DISP1) we find that transition between ‘inward-open’ and solvent ‘occluded’ states is accompanied by Na^+^ induced pinching of intracellular helical segments. Thus, our findings illuminate the energetics and ion-coupling stoichiometries of PTCH1 transport mechanisms, whereby 1-3 Na^+^ or 2-3 K^+^ couple to cholesterol export, and provide the first molecular description of transitions between distinct transport states.

## Introduction

Discerning the direction, energetics and stoichiometries of substrate transport by membrane proteins is essential for a holistic understanding of their molecular functions. Patched1 (PTCH1) is an integral membrane protein of the vertebrate Hedgehog (HH) signalling pathway which inhibits the Class F G protein-coupled receptor Smoothened (SMO) to prevent HH signalling^1^. PTCH1 inhibition of SMO is relieved by binding of Sonic Hedgehog (SHH) to the PTCH1 extracellular domain (ECD)^2^ (Fig. 1A, Supplementary Fig. 1A-B). HH pathway dysregulation results in severe developmental defects and compounded cancer pathologies^3^. Current FDA approved drugs targeting SMO are often overcome by tumorigenic resistance^4^, hence alternative avenues for pharmaceutical intervention must be explored.

**Figure 1:**
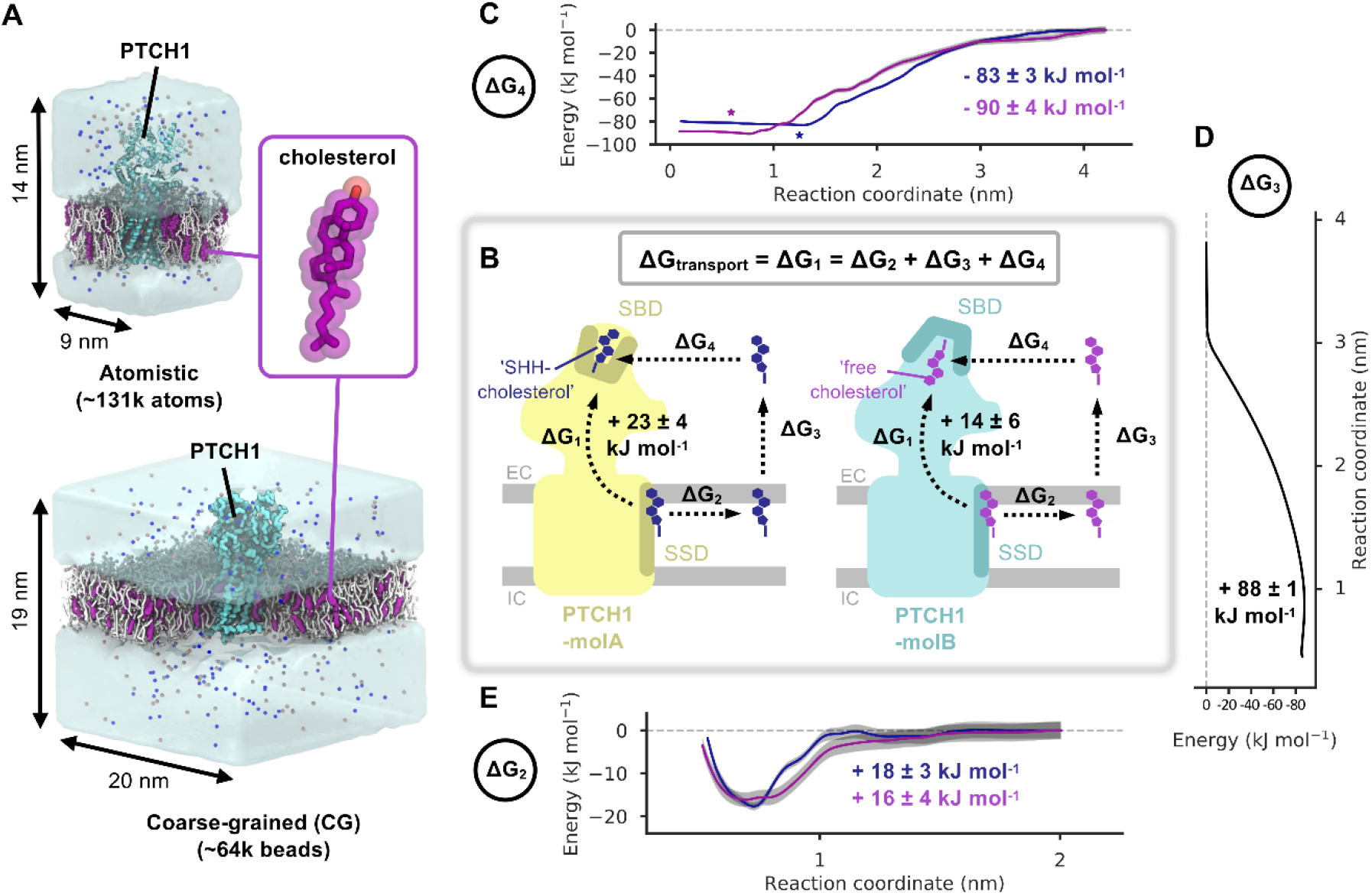
Patched1 and the energetics of cholesterol transport - the indirect pathway. **A)** Coarsegrained (CG) and atomistic simulation setups of PTCH1 (light blue, PDB: 6RVD^15^ (CG)/6DMY^5^ (atomistic)) embedded in 3:1 POPC:cholesterol (white/purple) bilayers. Water is shown as transparent surface and Na^+^/Cl^-^ ions are shown as blue/salmon spheres respectively. The inset shows the structure of cholesterol. **B)** Schematic diagram of the free energy changes associated with cholesterol movement between the PTCH1 sterol sensing domain (SSD) and the sterol binding domain (SBD) for the direct (ΔG_1_) and indirect (ΔG_2_, ΔG_3_, ΔG_4_) pathways. PTCH1-molA (yellow) and PTCH1-molB (light blue) are shown with either the ‘SHH-cholestero? (dark blue) or the ‘free cholesterol’ (purple) molecules positioned in the SBD and SSD (teal/ochre). The position of the extracellular (EC) and intracellular (IC) membrane leaflets are indicated in grey. **C-E)** CG potential of mean force (PMF) profiles for movement of ‘free cholesterol’ (purple) and ‘SHH-cholesterol’ (dark blue) between the SBD and the solvent (ΔG_4_) (**C**) or the SSD and the bulk membrane (ΔG_2_) (**E**) or for extraction of cholesterol from the membrane into the solvent (ΔG_3_) (**D**). Bayesian bootstrapping (2000 rounds) was used to estimate profile errors (grey). Stars indicate the position of the ‘SHH-cholesterol’ (within PTCH1-molA) and ‘free cholesterol’ (within PTCH1-molB) densities in a revised cryo-EM structure (PDB: 6RVD^15^).

The exact mechanism of PTCH1 mediated SMO inhibition remains ambiguous, despite decades of biochemical and cellular studies^1^, and a handful of recent PTCH1 structures^5–13^. Current models suggest PTCH1 may function as a cholesterol transporter to modulate the abundance of accessible membrane cholesterol available to bind and activate SMO^14^. Support for the proposed PTCH1 transport function is provided by observations of sterol-like densities within the ECD of PTCH1^5,8–10,15^ (Supplementary Fig. 1C). SHH inhibits PTCH1 by inserting a C-terminally linked cholesterol moiety into the PTCH1 ECD (Supplementary Fig. 1), blocking a putative cholesterol transport tunnel^15^. PTCH1 also shares homology with members of the prokaryotic resistance-nodulation-cell division (RND) family which export drugs, metal ions or small-molecules through their ECDs^16^. Additionally, PTCH1 is homologous to the Niemann-Pick disease type-C1 (NPC1) protein which transports LDL-derived cholesterol from the lysosomal lumen to the cytoplasm^17^. Thus, establishing the energetics of PTCH1 mediated cholesterol transport is needed to assess the feasibility of proposed transport models and further dissect how PTCH1 may inhibit SMO.

Classically, membrane transporters are characterised using vesicular uptake assays to assess the substrate specificities and ion-coupling requirements of transport. No such direct assay exists for PTCH1 function owing to a) difficulties in labelling the proposed sterol substrates, b) challenges with separating pre and post transport sterol pools (i.e. they may both reside within different microdomains of the same membrane)^18^, and c) problems with reconstituting functional PTCH1 owing to a requirement for CDO/BOC partner proteins or specific membrane topologies^19^. For example, PTCH1 localises at the base of the primary cilia in a highly curved invaginated membrane called the ciliary pocket^20^. Consequently, PTCH1 activity is usually assayed *in vivo* via downstream readouts such as HH target gene mRNA levels^21^, SMO localisation to the cilia^22^ or secondary labelling with e.g. PFO-based probes for distinct cholesterol pools^9,18^. Thus, there is a clear need for systematic investigation of PTCH1 function at the single molecule level in parallel with cellular readouts.

Molecular dynamics (MD) simulations enable the inspection of protein dynamics and interactions at atomic and near-atomistic resolutions (e.g. via coarse-grained (CG) methods)^23,24^. Increasingly, combinatorial MD and free energy methods are used to assess whether protein interactions with the surrounding environment are energetically favourable^25^. *In silico* methods can be used to obtain free energy values of ligands/lipids binding to proteins or transitioning between membrane and aqueous environments^26–28^. Previous MD simulations of HH pathway components have been used to e.g. demonstrate the stability of a novel cholesterol binding site in SMO^29^, dissect allosteric coupling between SMO cholesterol binding sites^30^ and explore PTCH1/SMO interactions with the membrane^25,31^.

Here we combine extensive MD simulations, free energy calculations, *in silico* mutants and *in vivo* biochemical assays of PTCH1 variants to explore cholesterol transport and ion-coupling by PTCH1. We find cholesterol export is associated with a positive free energy penalty, and would thus require coupling to an energetic input to be made favourable. We observe binding of Na^+^/K^+^ cations to a conserved site within the PTCH1 transmembrane domain (TMD) with similar affinity, which could provide the energetic impetus for cholesterol transport. Concurrently, we use simulations of Dispatched1 (DISP1), a HH pathway protein that utilizes the cellular Na^+^ gradient to release lipidated SHH from the membrane, to identify a mechanism of cation driven conformational changes via wetting of an intracellular cavity. The combined simulations, totalling ~1.5 ms, provide the first energetic assessment of cholesterol transport through PTCH1, illuminating the directionality and ion coupling stoichiometries of PTCH1 transport and ion driven conformational changes within the conserved RND family TMD.

## Materials and Methods

### Coarse-grained potential of mean force calculations

We previously performed CG MD simulations of PTCH1-molA and PTCH1-molB from the re-built PTCH1-SHH (2:1) structure (Protein Data Bank (PDB): 6RVD) embedded in symmetric POPC:CHOL (3:1) bilayers^15,32^(Fig. 1A). Snapshots from one 10 μs simulation of PTCH1-molA and PTCH1-molB were selected for use in potential of mean force (PMF) calculations. In total, nine PMFs were run for five coordinates (see Supplementary Table 1). PMF-1a used full length PTCH1-molA (with cholesterol bound to the sterol binding domain (SBD) in the ‘SHH-cholesterol’ orientation) or PTCH1-molB (with cholesterol bound in the ‘free cholesterol’ orientation). PMF-1b was built from the final window of PMF-1a with the cholesterols positioned at the base of the ECD. PMF-4 used the PTCH1-molA and PTCH1-molB ECDs with the ‘SHH-cholesterol’ and ‘free cholesterol’ bound to the SBD as in PMF-1a (and according to the cryo-EM densities). PMF-3 used a 10 x 10 nm^2^ bilayer patch (in the absence of protein). PMF-2 was constructed as in PMF-1a but with cholesterol bound to the sterol sensing domain (SSD) instead of the SBD. See Extended Methods for full system details including setup and cholesterol orientations.

PMF calculations were carried out in accordance with a previously published method^33^. GROMACS 2018 and 2019 (www.gromacs.org) were used to perform all simulations^34^. Steered MD simulations were used to pull the cholesterol of interest through the protein (PMF-1), away from the protein (PMF-2, PMF-4) or out of the membrane (PMF-3). These followed a 1D reaction coordinate between the centre of mass (COM) of cholesterol and a backbone bead of PTCH1 (PMF-1, −2 and −4, see Extended Methods) or the COM of the bilayer (PMF-3). An umbrella pulling force of 1000 kJ mol^-1^ nm^-2^ and a pulling rate of 0.1 nm ns^-1^ were used in all cases with the exception of PMF-1b where the pulling rate was 10 nm ns^-1^ to prevent extraneous pulling of cholesterol into the aqueous solvent after egression from the ECD. Windows were spaced every 0.05 nm along the reaction coordinate and simulated for 1-3 μs each until convergence (Supplementary Table 1 and Supplementary Fig. 2). A 1000 kJ mol^-1^ nm^-2^ umbrella potential was used to limit cholesterol movement within each window. PMF profiles were constructed using the weighted histogram analysis method (WHAM) implemented in *gmx wham*^35,36^. The first 200 ns of each window was discarded as equilibration time and 2000 Bayesian bootstraps were used to estimate the PMF error. Since PMFs 2-4 are independent, the total error of the indirect pathway was calculated as in Equation 1. Setup and analysis of PMFs was aided by use of the *pmf.py* tool (DOI:10.5281/zenodo.3592318)^33^.

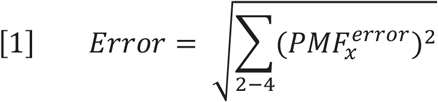

### Coarse-grained absolute binding free energy calculations

Absolute binding free energy (ABFE) calculations^37^ were performed on the CG PTCH1 complex with a cholesterol molecule bound to the TMD SSD (just PTCH1-molA), the ECD SBD and a mid-point along PMF-1a at approximately 2.4. nm on the reaction coordinate (for PTCH1-molA and PTCH1-molB). Initial poses reflect those used to seed the steered MD simulations as described above. In all cases, the systems were trimmed to *ca.* 15 x 15 x 15 nm^3^ and for the TMD and mid-point systems all cholesterol apart from the target molecule were removed from the system, followed by 200 ns of equilibration.

ABFE calculations were essentially run as previously described^33,38^ (Supplementary Table 2). In brief, the target cholesterol is fully decoupled from the system along an alchemical coordinate, λ. The Lennard-Jones interactions of the cholesterol particles were switched off over 29 windows, using a step size of 0.05 for λ=0-0.6 and a step size of 0.25 for λ=0.6-1. A soft-core potential was used, with an α of 0.5 and a σ of 0.3. For each window, the systems were minimised using steepest descents before 3 x 250 ns production simulations, using a stochastic integrator at 323 K, with pressure held at 1 bar using the Parrinello-Rahman barostat^39^, with a *τ*_p_ of 4 ps and a compressibility of 5 x 10^-5^ bar^-1^. Throughout the λ windows, flat-bottomed distance restraints of 10 kJ mol^-1^ nm^-2^ were applied between the cholesterol and its bound state to keep it in the binding site at high values of λ^40,41^. The bound state was defined using two dummy beads whose position updated in relation to reference beads in the binding site: E274+S331 in the SBD site for both states, A936+Y1013/Y801 in the mid-point site for PTCH1-molA/B, and A478+V481 for the SSD pose. At low values of λ, the binding site itself restricts the cholesterol molecule, and the restraints have a negligible effect. The computed energies were constructed into energy landscapes along λ using Multistate Bennett Acceptance Ratio (MBAR)^42^ as implemented in alchemical analysis^43^.

### Atomistic simulations of PTCH1

The cryo-EM structure of PTCH1 with a putative ion density visible in the TMD (PDB: 6DMY)^5^ was selected for use in atomistic simulations (Fig. 1A). PTCH1 was setup and embedded within a POPC:CHOL (3:1) bilayer as described in the Extended Methods. A Na^+^ ion was positioned within the TMD according to the cryo-EM density^5,15^ and the system was solvated with TIP4P water and approximately 0.15 M NaCl. Two rounds of steepest decent energy minimisation were proceeded by 2 x 5 ns NVT and NPT equilibration steps with restraints applied to PTCH1.

Atomistic simulations (3 x 100 ns) were performed with one Na^+^ initially bound in the TMD. Additional 3 x 100 ns simulations were run in the presence of −100 mV and −200 mV membrane voltages. 3 x 50 ns simulations were also run without Na^+^ bound and 3 x 100 ns simulations were performed in the presence of 0.15 KCl in replace of NaCl. 3 x 100 ns were initiated with either 3 x Na^+^ or 3 x K^+^ ions bound at equivalent positions to in the DISP1 structure (PDB: 7RPH)^44^. *In silico* mutants of PTCH1 were each simulated for 3 x 50 ns (Extended Methods, Supplementary Table 3).

GROMACS 2018 and 2019 (www.gromacs.org) were used to run simulations^34^. A 2 fs timestep was used and the CHARMM36 forcefield^45^ described all components. The Nosé-Hoover thermostat was used to maintain temperature at 310 K (*τ*_t_=0.5 ps)^46,47^. The Parrinello-Rahman barostat^39^ was used to maintain semi-isotropic pressure at 1 bar (*τ*_p_=2.0 ps), with a compressibility of 4.5×10^-5^ bar^-1^. The PME method was used to model long-range electrostatic interactions^48^. Van der Waals interactions were cut off at 1.2 nm. Bond lengths were constrained to the equilibrium values using the LINCS algorithm^49^. A dispersion correction was not applied.

### Atomistic simulations of DISP1

A DISP1 structure bound to three Na^+^ ions was used in atomistic simulations (PDB: 7RPH, ‘R conformation’)^44^. DISP1 was setup and simulated as described in the Extended Methods in distinct ion bound or apo states, 3 of which were generated by sequential ion removal from the final frame of the previous trajectory (3 x 100 ns x 5 ion bound states) (Supplementary Table 3). Simulation parameters were identical to those described above for PTCH1.

### Cation free energy perturbation

The end snapshot from one 100 ns atomistic simulations of PTCH1 with Na^+^ bound in the TMD was used in free energy perturbation (FEP) calculations^43^. Na^+^ was alchemically changed to K^+^ over 31 windows by independently modifying the mass (λ windows 0-10) and VDW radii (λ windows 11-30) of the bound ion using a dual topology approach. Mass modification did not contribute significantly to the overall free energy and therefore a λ step size of 0.1 was used in windows 0-10 and a step size of 0.05 in windows 11-30. A soft-core parameter (σ = 0.3) was used for Lennard-Jones interactions. Each window was simulated for 50 ns using the protocol described above for atomistic simulations of PTCH1 with the exception that a stochastic dynamics integrator was used. Perturbation of Na^+^ to K^+^ in free solution was achieved using a single Na^+^ in a 3 x 3 x 3 nm^3^ TIP4P water box. A single Cl^-^ counter ion was present. The solvent box was equilibrated for 20 ns before use in FEP calculations to allow water to equilibrate around the ion. Each window was simulated for 5 ns using the same protocol described for the PTCH1 bound Na^+^ systems. 5 repeats of each FEP calculation were performed. 5 repeats of the inverse perturbations (K^+^ to Na^+^) in the bound and free states were also performed which gave free energy values of the same magnitude but inverse signs (Supplementary Fig. 3). Alchemical analysis^43^ was used in calculation of the overall free energy values from λ windows. There was good agreement between analysis methods and so values calculated using MBAR^42^ are reported as the free energy mean and standard deviation between repeats (Supplementary Fig. 4). The first 200 ps of each window were discarded to allow for equilibration.

### Generation of stable cell lines expressing Patched1 mutants

#### Patched1 construct generation

Human Patched1 (hPtch1), comprising amino acids 75-1185 including a deletion of intracellular loop 3 (ICL3) (Δ630–717), was cloned into the pHR-CMV-TetO2 vector^50^ (Addgene plasmid #113893). For functional studies the following single or double point mutations were introduced: V510F, D513Y, L517C/P1125C, L570C/V1081C, and P115R/N802E.

#### Virus generation

Viral supernatant containing the hPtch1 variants was generated using HEK 293T cells (ATCC; Cat#CRL03216) and then added to *Ptch1* knockout mouse embryonic fibroblasts (*Ptch1^-/-^* MEFs)^51^. To generate virus, 2.5 x 10^6^ HEK 293T cells were plated in 6 cm plates in 3 mL of antibiotic-free High Glucose DMEM supplemented with L-glutamine, sodium pyruvate, and non-essential amino acids. Twenty-four hours later, cells were transfected by mixing 500 μL room temperature Optimem, 5 μg of the pHR-CMV-TetO2 plasmid containing the hPtch1 gene, 2.5 μg pVSVG, 3.75 μg psPAX2, and 45 μL PEI (4 μL of PEI to 1 μg of DNA). After incubating this mixture at room temperature for 20 minutes, it was added dropwise to the 6 cm dish of HEK 293T cells. The following day, media on the 293T plates was exchanged for fresh media. At this time, 6 cm plates were seeded with 2 x 10^5^ *Ptch1^-/-^* MEFs. Twenty-four hours later, the virus-containing media was harvested from the 293T cells, and polybrene was added to 4 μg/mL. This undiluted viral supernatant was added directly to the *Ptch1^-/-^* MEFs.· Fresh antibiotic-free DMEM was added to the 293T cells for a second round of virus production, followed by a second infection of the *Ptch1^-/-^* MEFs 24 hours later.

#### Isolation of Ptch1-expressing cells and validation

*Ptch1^-/-^* MEFS containing the integrated hPtch1 variants were trypsinized and analysed compared to an un-infected (control) population using Fluorescence Activated Cell Sorting (FACS). A population of cells with fluorescence from the integrated hPtch1-mVenus-1D4 construct was clearly visible. This population was used to set a sorting gate for the cells containing hPtch1 gene. Approximately 250,000 cells were sorted and then plated into a 6 cm dish. Once these cells were grown to a sufficient density, they were seeded into additional plates for Western blot analysis^52^ to confirm expression of the hPTCH1-mVenus-1D4 gene.

## Results

### Energetic barrier for cholesterol export through PTCH1

We constructed free energy cycles corresponding to cholesterol movement between the transmembrane and extracellular domains of PTCH1-molA and PTCH1-molB (Fig. 1). For PTCH1-molA and PTCH1-molB, each cycle is composed of a pathway directly linking the sterol sensing domain (SSD) and sterol binding domain (SBD) through PTCH1 (ΔG_transport_ = ΔG_1_), and an indirect pathway between the same sites (composed of ΔG_2_, ΔG_3_ and ΔG_4_) via the surrounding environment (Fig. 1B). Thus, free energy cycles were constructed to compare how the energetic landscape changes between the PTCH1-molA and PTCH1-molB conformations and between the ‘free cholesterol’ and ‘SHH-cholesterol’ orientations which differ by ~180°.

#### The indirect pathway

We used potential of mean force (PMF) calculations with umbrella-sampling and a CG force-field^53^ to compute the indirect pathway free energy calculations. The indirect pathway was modelled using three PMFs corresponding to movement of cholesterol between the PTCH1 SSD and the bulk membrane (PMF-2: ΔG_2_), cholesterol movement between the membrane and solvent (PMF-3: ΔG_3_) and movement of cholesterol between the solvent and the SBD of PTCH1 (PMF-4: ΔG_4_). In PMF-4, the SBD provides a favourable environment for cholesterol binding relative to the solvent in both PTCH1 conformations (Fig. 1C, PTCH1-molA: ΔG_4_ = −83 ± 3 kJ mol^-1^, PTCH1-molB: ΔG_4_ = −90 ± 4 kJ mol^-1^). The PMF-4 landscape within the PTCH1 SBD is flat suggesting cholesterol may slide along the tunnel-like cavity within the SBD without significant energetic penalty as opposed to localising within a well-defined binding site. This may allow cholesterol to deviate from the cryo-EM observed SBD binding poses (starred in Fig. 1C). For PMF-3, extraction of cholesterol from its favourable hydrophobic bilayer environment into the surrounding solvent was associated with a large, positive free energy change (Fig. 1D, ΔG_3_ = +88 ± 1 kJ mol^-1^) comparable to previous reports^54,55^. Indeed, the magnitude of the ΔG values for PMF-4 and PMF-3 compare well, emphasising the hydrophobicity of the PTCH1 SBD and the selection pressure on PTCH1 to stabilise cholesterol in a domain that mimics cholesterol stabilisation in its native environment.

The resolution within the detergent region of existing cryo-EM PTCH1 structures limits the unambiguous assignment of sterol densities which has led to uncertainty in modelling the orientation of cholesterol within the SSD^5–11^. For this reason, in PMF-2 we compared the free energy change of cholesterol movement out of the PTCH1 SSD when the ROH bead (equivalent to the 3β-hydroxyl (3β-OH) group of cholesterol in atomistic resolution) pointed towards the extracellular leaflet headgroups (‘OH-up’) or towards the bilayer midplane (‘OH-down’). For the ‘OH-up’ orientation we observe a well-defined binding site with a moderate free energy change (Fig. 1E, PTCH1-molA: ΔG_2_ = +18 ± 3 kJ mol^-1^, PTCH1-molB: +16 ± 4). By contrast, there was no observable well for cholesterol binding to PTCH1 SSD in the ‘OH-down’ orientation and the profile remained approximately flat over the reaction coordinate (Supplementary Fig. 5, PTCH1-molB: +4 ± 3 kJ mol^-1^). From this we conclude that the most likely cholesterol orientation within the PTCH1 SSD is the ‘OH-up’ configuration (as reported in Fig. 1E), consistent with density modelled in the refined cryo-EM structure^15^.

Using the free energy values from the indirect pathway we obtained an overall ΔG value for cholesterol movement from the PTCH1 SSD to the SBD (ΔG_1_). For PTCH1-molA and PTCH1-molB we calculate ΔG_1_ to be +23 ± 4 and +14 ± 6 kJ mol^-1^ respectively, i.e. cholesterol binding to the SSD within the TMD is substantially more favourable than binding to the extracellular SBD. Thus, if PTCH1 functions as a cholesterol exporter, an energetic input would likely be required for the cholesterol to move from the SSD to the SBD. Similarly, the above data are also consistent with spontaneous movement of cholesterol from the PTCH1 SBD to the SSD membrane binding site, providing that a pathway is available.

#### The direct pathway

Previous atomistic simulations and analysis of PTCH1 structures identified putative sterol transport tunnels throughout the PTCH1 ECD^5,7,11,15^. These tunnels form between the PTCH1 SBD and cavities within the ECD in proximity to either the SSD or TM12 at the membrane interface, or side exits between PTCH1 ECD1 and ECD2. In contrast, the pathway for sterol movement between the membrane-proximal region of the PTCH1 ECD to the SSD binding site on the PTCH1 TMD is uncertain. Indeed, current predictions have not yielded a consensus pathway^5,7,15^. Thus, while it is possible to form a PMF pathway between the SBD and the base of the ECD, using the previously identified tunnels to guide selection of an appropriate reaction coordinate, elucidation of a plausible pathway between the ECD base and the SSD transmembrane site is non-trivial. For this reason, we used a combination of absolute binding free energy (ABFE) calculations^37^ to directly probe the free energy difference between cholesterol bound to the SBD and the SSD sites, and PMF calculations confined between the SBD and ECD base to understand the energetic landscape over this subsection of the direct pathway (Fig. 2A).

**Figure 2:**
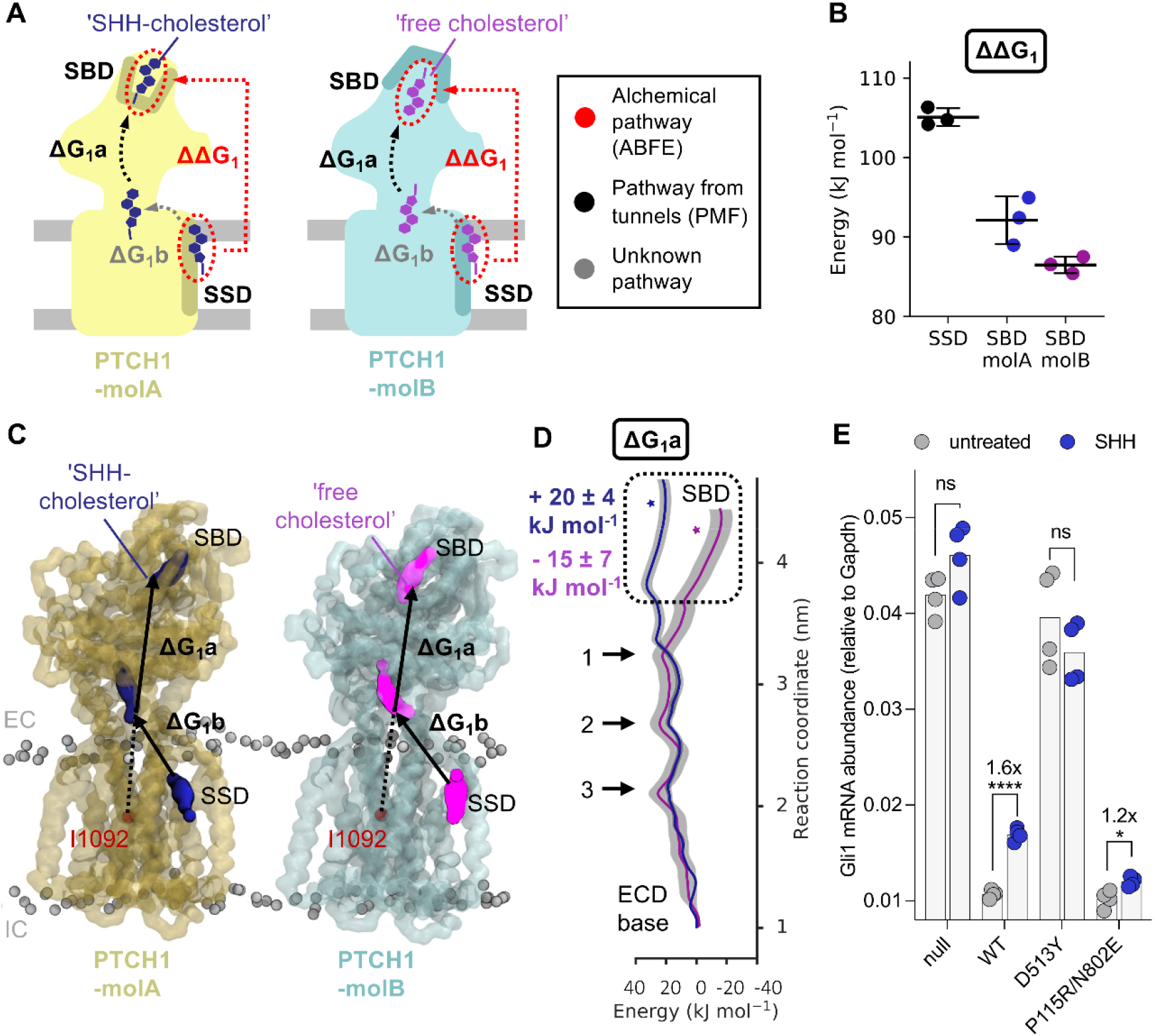
Cholesterol transport energetics - the direct pathway. **A)** Schematic diagram of the free energy changes associated with cholesterol movement between the SSD and SBD of PTCH1 for the direct pathway, coloured identically to in Fig. 1B. Decoupling of cholesterol from the SBD/SSD sites are indicated by red circles, with the difference between them (ΔΔG_1_) shown as a red arrow to indicate alchemical transformation. A black arrow indicates shows movement of cholesterol between the ECD base and the SBD, used in PMF calculations to derive ΔG_1_a. Grey arrows correspond to currently uncertain regions of the transport pathway. **B)** The free energy values of decoupling cholesterol from the SSD (PTCH1-molA) and SBD (PTCH1-molA and PTCH1-molB) as obtained from absolute binding free energy (ABFE) calculations. **C)** Snapshots from PMF-1a indicating movement of cholesterol between the ECD base and SBD of PTCH1 (residues used in the steered MD are labelled in red). CG representation of PTCH1 backbone beads are shown in transparent light blue/yellow, ‘free cholesterol’ is coloured purple, ‘SHH-cholesterol’ is coloured dark blue and lipid phosphate beads are shown in grey. **D)** PMF profile for cholesterol movement through the ECD of PTCH1-molA/B (ΔG_1_a) (see **C**). Bootstrapping errors (2000 rounds) are shown in grey. The position of cholesterol within the SBD pocket in the cryo-EM model (PDB: 6RVD^15^) is starred. Arrows in **D** indicate energetic peaks 1-3 within the ECD core conserved between PTCH1-molA and PTCH1-molB (see Supplementary Fig. 7). **E)** HH signalling strength is determined by measuring endogenous *Gli1* mRNA abundances (normalized to the control *Gapdh*) in response to SHH ligands (200 nM, 20 hours) in *Ptch1^-/-^* cells stably expressing the indicated variants. Statistical significance is determined by a Student’s t-test with a Welch’s correction. Exact *P* values for comparisons: *Ptch1^-/-^* untreated vs. SHH = 0.085, WT untreated vs. SHH < 0.0001, D513Y untreated vs. SHH = 0.2712, and P155R/N802E untreated vs. SHH = 0.0205. not significant (ns), \**P* > 0.05, \**P* ≤ 0.05, \*\**P* ≤ 0.01, \*\*\**P* ≤ 0.001, and *\*\*\*\*P* ≤ 0.0001.

To obtain a second estimate of ΔG_1_ we applied CG ABFE calculations to directly probe the free energy difference between the ‘SHH-cholesterol’ (in PTCH1-molA) and ‘free cholesterol’ (in PTCH1-molB) by alchemically decoupling cholesterol from the SSD and SBD of each PTCH1 conformation (Fig. 2B). The difference between decoupling from the SSD and SBD sites can be used to obtain an estimate of ΔG_1_ (or ΔΔG_1_ as is the case here, where ΔΔG_1_ = ΔG_1-SSD_ – ΔG_1-SBD_) that does not rely on formation of a physical pathway through PTCH1 (Fig. 2A). Our ABFE calculations predict a positive free energy change between the SSD and SBD sites for both PTCH1 molecules via the direct alchemical pathway, i.e. cholesterol binding to the SSD is more favourable than the SBD (Fig. 2B, PTCH1-molA: ΔΔG_1_ = +13 ± 3 kJ mol^-1^, PTCH1-molB: ΔΔG_1_ = +20 ± 1 kJ mol^-1^). While the magnitude of the direct pathway free energy change is reversed for PTCH1-molA and PTCH1-molB compared to the indirect pathway calculations, these data are consistent with a predicted increase in free energy between the SSD and SBD sites (Fig. 3) and therefore support the proposed hypothesis that coupling to an energy source would be required for PTCH1 to export cholesterol.

**Figure 3:**
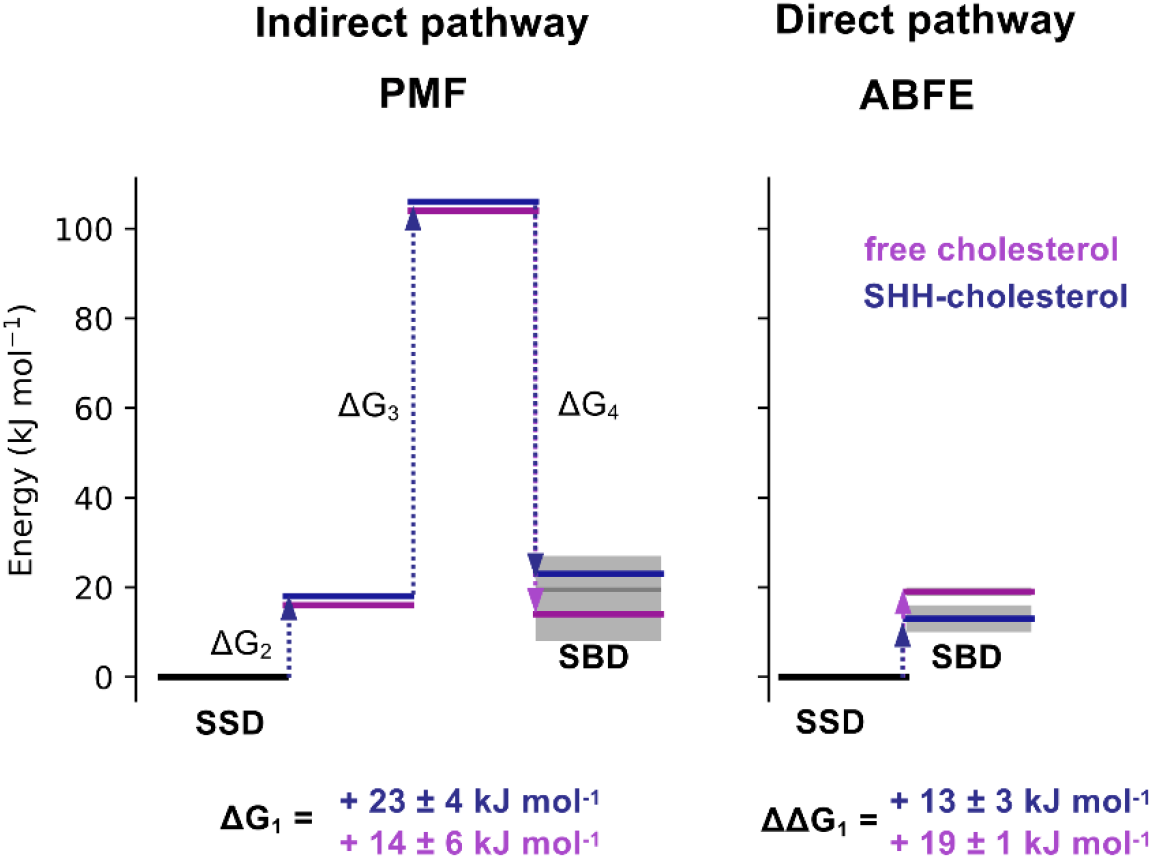
Movement of cholesterol between the SSD and SBD is associated with an energetic barrier. Comparison of ΔG_1_ values for ‘free cholesterol’ (purple) and ‘SHH-cholesterol’ (dark blue) movement between the PTCH1 SSD and SBD, derived from the indirect (PMF, Fig. 1) and direct (ABFE, Fig. 2) pathways. Step wise free energy changes are shown as connecting arrows and are labelled. ΔG_1_ from the indirect pathway was calculated in quadrature (Equation 1) since PMFs 2-4 were independent. In all cases cholesterol movement from the SSD to the SBD gives ΔG > 0 kJ mol^-1^.

We performed PMFs of moving cholesterol between the SBD and the base of the ECD (PMF-1a: ΔG_1_a) to better understand the energetic landscape over the most well-defined bisection of the proposed sterol transport tunnel (Fig. 2C, Supplementary Fig. 6A). PTCH1-molA and PTCH1-molB show a similar energetic profile for cholesterol movement through the lower and mid regions of the ECD, consisting of three peaks of approximately +18 to +27 kJ mol^-1^ (Fig. 2D). This is suggestive of a ‘pez-like’ transport mechanism formed by multiple, similar binding sites within the ECD. Overlay of the profiles in this region was confirmed by performing ABFE calculations of the two cholesterol molecules bound within a central portion of the ECD (2.4 nm on PMF-1a) which yielded similar ΔG values (PTCH1-molA: + 50 ± 1 kJ mol^-1^, PTCH1-molB: + 55 ± 2 kJ mol^-1^). Strikingly, as cholesterol moves towards the SBD the paths for PTCH1-molA and PTCH1-molB diverge. Cholesterol movement towards the cryo-EM binding pose within the SBD of PTCH1-molB (purple star in Fig. 2D) is associated with a stabilisation of −15 ± 7 kJ mol^-1^ compared to the free energy at the ECD base (1 nm in Fig. 2D). Conversely, movement of cholesterol towards the PTCH1-molA SBD pose (dark blue star in Fig. 2D) is associated with an increase in free energy of +20 ± 4 kJ mol^-1^. This is consistent with previous atomistic simulations which show a stabilisation of the ECD upper lobe by cholesterol in PTCH1-molB but not PTCH1-molA, as demonstrated by a reduced RMSD compared to the apo ECD conformations^15^. This result also corroborates our PMF-4 profiles which suggest the PTCH1-molB SBD cholesterol binding site is stabilised relative to PTCH1-molA, albeit to a lesser extent. Further, in all PTCH1 structures solved to date with a SBD conformation matching that of PTCH1-molB, sterol like density is observed in an equivalent region to the ‘free cholesterol’ molecule (Supplementary Fig. 1C)^5,8–10^. Together with our observations of a free energy well towards the PTCH1-molB SBD, this suggests the ‘free cholesterol’ site may represent a local binding site within the ECD, perhaps facilitating downhill (i.e. favourable) movement of cholesterol towards the SBD to promote transport directionality.

Due to the uncertainties in the transport pathway between the SSD and the ECD base (described above) it is not possible to obtain full characterisation of the energetic landscape using PMF calculations over the membrane proximal region of PTCH1. This is complicated by the requirement for the ‘free cholesterol’ to flip between the ECD base and SSD sites in PTCH1-molB, potentiating difficulties with obtaining this PMF (Fig. 2A). Lipid re-orientation is also observed in other RND proteins such as in MmpL3^56^. Our attempts (PMF-1b: ΔG_1_b), confined to Supplementary Fig. 6B-C, show a large energetic barrier which occupies the central region of the PMF-1b profile. This occurs as a result of solvation of the hydrophobic cholesterol molecule as it leaves the tunnel cavity at the base of the ECD and slides along the protein surface towards the bilayer. It seems unlikely that this represents a physiological pathway considering the efficiency of HH signalling pathway inhibition by PTCH1 and the high insolubility of cholesterol and, as such, is excluded from our predictions of ΔG_1_.

In summary, our ΔG_1_ values calculated from the indirect (PMF) and direct (ABFE) pathways both suggest moderate energetic costs for cholesterol movement from the SSD to the SBD of approximately +13 to +23 kJ mol^-1^ (Fig. 3).

### Blocking the ECD sterol tunnel does not disrupt PTCH1 function

We sought to identify those residues which may contribute to energetic bottlenecks in the cholesterol pathway through PTCH1 ECD and which may therefore provide insight into plausible therapeutic interventions. We identified residues within 0.6 nm of cholesterol in those windows that surround energetic peaks in the PMF-1a profile (Fig. 2D, see marked arrows). The four residues with the highest occupancy i.e. residues which are within 0.6 nm of cholesterol for the greatest fraction of the simulation time were mapped onto the structure of PTCH1 at each peak (Supplementary Fig. 7A). The residues localise to two layers at the ECD1-ECD2 interface which we term the upper and lower restrictions. Visualisation of the PMF window trajectories suggests cholesterol interaction with the upper and lower restrictions results in the first and third energetic peaks respectively whereas the middle peak is formed by cholesterol interacting with both the lower and upper restrictions simultaneously. We also calculated the per residue cholesterol occupancies over all PMF-1a windows which showed residues comprising the upper and lower restrictions also had the highest occupancies over the whole PMF-1a profile (Supplementary Fig. 7B). There is good agreement between identified residues from PTCH1-molA and PTCH1-molB due to conservation of the energetic profile over the lower and mid regions of the ECD (Fig. 2D). In particular, in both PTCH1 molecules L227, M335, W337, A936 and Q938 contribute to the upper restriction just below the base of the SBD pocket, and P155 and N802 form part of the lower restriction just above a previously identified sterol site^9^ near the entrance to the putative sterol conduit at the base of the ECD.

Observed sterol densities in PTCH1 structures do not necessarily indicate cholesterol movement between these sites (or within the internal PTCH1 ECD cavity we probed computationally). Sterol detergents added during protein purification (e.g. CHS) may bind to distinct sites without interconnected sterol translocation through the ECD cavity. Given the +18-27 kJ mol^-1^ energetic penalty associated with cholesterol movement through the lower and upper restrictions (Fig. 2D), we sought to assess whether cholesterol movement through the ECD is necessary for PTCH1 function in cells. To further dissect whether proposed sites represent distinct cholesterol binding sites or a genuine interconnected cholesterol pathway we generated a salt-bridge mutant (P155R/N802E) which occludes the proposed PTCH1 ECD transport cavity (Fig. 2E, Supplementary Fig. 8). We generated cell lines stably expressing hPTCH1 variants in a *Ptch1* knockout background to cleanly assess their function in HH signalling activity assays. As expected, cells lacking *Ptch1* (*Ptch1^-/-^*) show high basal HH strength due to unrestrained SMO signalling downstream (Fig. 2E). Expression of either wildtype (WT) hPTCH1 or the P155R/N802E mutant blocked SMO activity, reducing HH signalling strength. As a control, we expressed an inactive PTCH1 mutant, D513Y, which was unable to block HH signalling in the *Ptch1^-/-^* background. This result suggests that either a) introducing a salt-bridge to the cholesterol tunnel in PTCH1 does not block its function or b) this mutation is not sufficient to block cholesterol movement through the tunnel. Interestingly, the fold-change in pathway activation upon SHH addition is reduced for P155R/N802E compared with cells expressing WT PTCH1. This suggests that the predicted salt bridge indeed forms and disrupts SHH binding, without disrupting PTCH1 activity. Hence, we provide cellular evidence that sterol translocation through the ECD may not be required for PTCH1 function, in agreement with previous observations that PTCH1 can still function to inhibit SMO in the absence of the whole ECD2 domain^51,57^.

### Identification of multiple, interconnected cation binding sites within PTCH1 TMD

PTCH1 shares a conserved structural architecture with members of the prokaryotic RND transporter superfamily, which utilise the proton gradient to export small molecules, drugs, peptides and metals^16,58^. The proposed RND proton binding site is formed by two or three anionic residues in the centre of the TMD^59^. PTCH1 sequence alignment has identified two aspartate residues (D513 and D514) on TM4 and a glutamate residue on TM10 (E1095) as forming part of highly conserved GXXXDD and GXXX(E/D) motifs at the centre of the TMD, in equivalent positions to the proton binding sites in RND transporters^5,9^.

We previously identified an ion-like density in proximity to the anionic triad within an existing PTCH1 structure (Fig. 4A, inset)^5,15^. Given our finding that cholesterol movement between the SSD and SBD is associated with a free energy barrier, we reasoned cholesterol export could be made favourable by coupling to cation binding and/or movement across the membrane.

**Figure 4:**
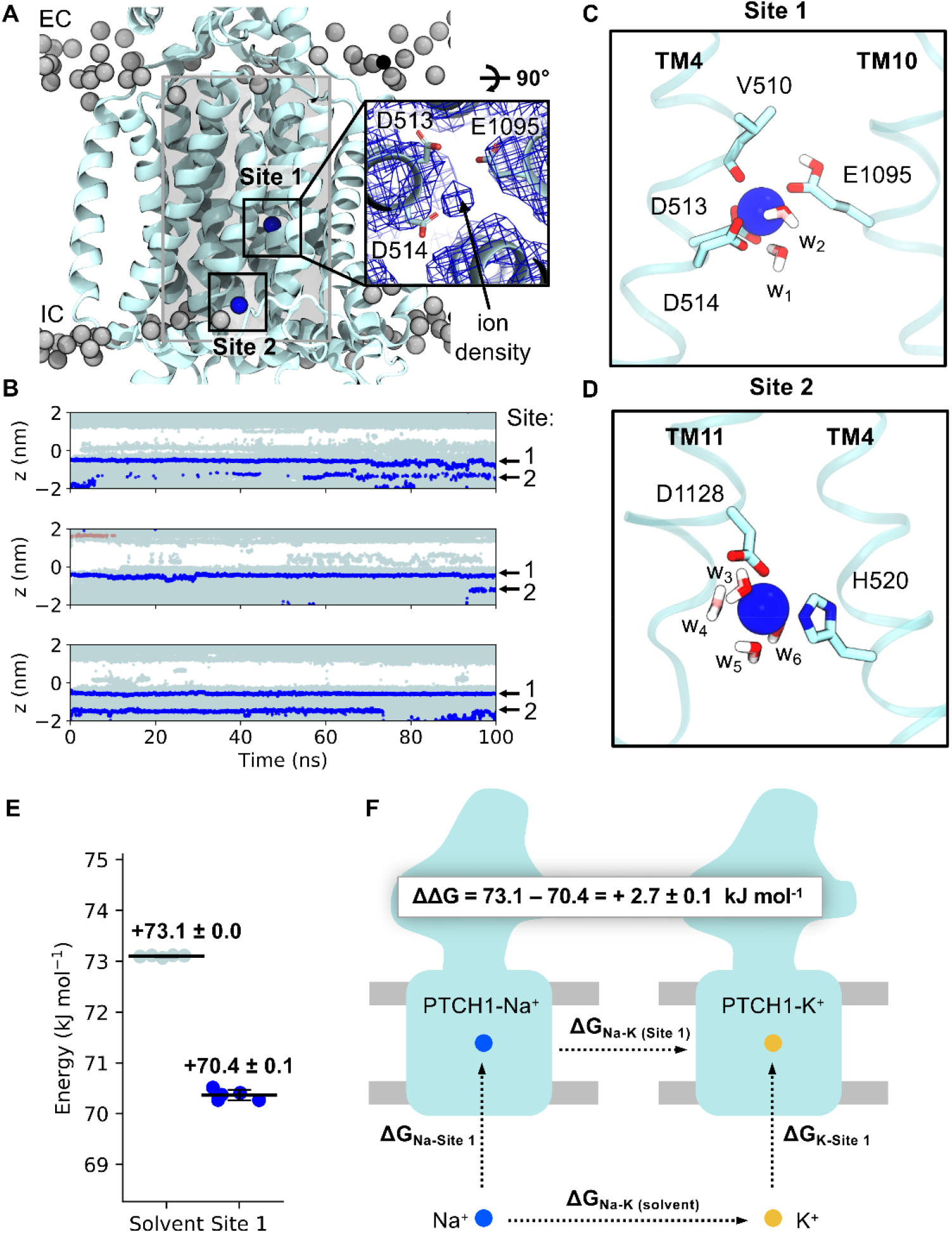
Identification and selectivity of cation binding sites within the PTCH1 TMD. **A)** Snapshot from atomistic simulations of PTCH1 (light blue, PDB: 6DMY^5^) indicating the location of cation binding sites within the TMD. Na^+^ ions are shown as blue spheres and lipid phosphates are grey. Inset: cation-like density surrounded by anionic tired residues at Site 1 within the cryo-EM structure^5^, as viewed from the extracellular face. A cylinder (length 4 nm, radius 1.3 nm) centred on the midpoint of V520 and I1092 Cα atoms is shown in grey and was used to identify water and ions within the PTCH1 TMD (**B)**. **B)** The z coordinates of water oxygen atoms (light blue), Na^+^ (blue) and Cl- (salmon) ions localised within the PTCH1 TMD (see **A**, grey cylinder) over the length of 3 x 100 ns simulations initiated with Na^+^ bound at the density observed in **A**. Snapshots of Site 1 (**C**) and Site 2 (**D**) showing coordination of bound Na^+^ ions by surrounding residues (stick representation) or waters (w1-6). **E)** Free energy perturbation (FEP) calculations for alchemical transformation of Na^+^ into K^+^ within the solvent or bound to Site 1. **F)** Schematic representation of the free energy cycle used to calculate the difference in Na^+^ binding to Site 1 compared to K^+^ (ΔΔG).

To test the viability of the proposed cation binding site we performed atomistic simulations of PTCH1 with Na^+^ located in the TMD according to the cryo-EM density at a position we term Site 1 (Fig. 4). Na^+^ remained bound at Site 1 over the 100 ns of 3 repeat trajectories with a mean RMSD of 0.15 ± 0.02 nm compared to the initial ion position (Fig. 4B). There is active debate within the PTCH1 literature on the molecular identity of the proposed coupling cation; with both Na^+^ and K^+^ proposedly implicated in PTCH1 function or regulation^21,22,52^. Hence, we performed further simulations, initiated with an apo TMD (in 0.15 M NaCl or KCl) and observed spontaneous Na^+^/K^+^ binding to Site 1 in all replicates (Supplementary Fig. 9A). Na^+^/K^+^ were coordinated octahedrally by six oxygen atoms from the carboxyl groups of D513^-1^, D514^-1^ and E1095^0^ (where superscript indicates the modelled charge of the residue), the backbone carbonyl of V510 and two water molecules (Fig. 4C). Residue-ion contact mapping showed occasional displacement of the water at position w2 by the hydroxy side chain of T551 (Table 1). Application of a membrane voltage of physiological magnitude was not sufficient to promote displacement of Na^+^ from Site 1 (Supplementary Fig. 9B). A second cation binding site (Site 2, Fig. 4), not visible in the cryo-EM data, was observed in the simulation data on TM4/TM11 (Fig. 4D), with an ion from bulk solvent entering a small solvent accessible cavity formed by local bilayer deformation at the intracellular PTCH1 surface (also visible in MemProtMD^60^). Here, Na^+^/K^+^ is coordinated by 4 water molecules, and two residues from the TMD, H520^0^ and D1128^-1^. A similar solvated cavity was observed in simulations of the RND member AcrB whereby a continuous solvent pathway extends from this surface cavity into the TMD core^61,62^. In some simulations Na^+^/K^+^ was observed to move from Site 2 into Site 1 mediated by flipping of D1128 towards TM4 to handover Na^+^ to D513, even when Site 1 was already occupied. This suggests that Site 2 might act as an intermediate site between the solvent and Site 1, and that anionic triad residues are capable of coordinating at least 2 cations simultaneously.

**Table 1:** Residue contacts with Na^+^ bound at Site 1 across 3 x 100 ns simulations of PTCH1. Residues with < 1 % contact are excluded.

| Residue | No. frames within 0.3 nm of Na<sup>+</sup> (%) |
| --- | --- |
| D514 | 93.3 |
| D513 | 89.5 |
| V510 | 85.1 |
| E1095 | 81.8 |
| T551 | 5.8 |

The suggested involvement of K^+^ in PTCH1 function^21,52^ is surprising given an absence of known eukaryotic K^+^ coupled transporters and the theoretical closeness of the resting plasma membrane potential and the equilibrium K^+^ potential^52^. To explore this further we calculated the free energy associated with perturbation of Na^+^ to K^+^ at Site 1 compared to in bulk solution using FEP calculations (Fig. 4E-F, Supplementary Fig. 3). Perturbation of Na^+^ to K^+^ was a marginal 2.7 ± 0.1 kJ mol^-1^ more favourable when bound to Site 1 than in solution, suggesting Na^+^/K^+^ bind to Site 1 with very similar affinity. Thus, it is perhaps unsurprising that both ions have been implicated in PTCH1 function/regulation biochemically^21,52^.

### An intracellular open solvent cavity forms within the PTCH1 TMD

To gain insight into the role of the PTCH1 TMD we analysed the TMD localised water density across atomistic simulations of PTCH1. A hydrated cavity encompassing the anionic triad residues and opening to the bulk intracellular solvent was clearly visible from both the water density analysis and the z coordinates of water and ions over time (Fig. 5A, Fig. 4B, Supplementary Fig. 9). Water entered the cavity between TM4 and TM10 at the base of the TMD or via the water pocket surrounding Site 2 (Fig. 5A, see arrows). The extracellular TMD half remained de-wetted and water did not cross between the extracellular and intracellular compartments during the simulations, akin to previously observed water networks present in the O- and L-states of AcrB^61^.

**Figure 5:**
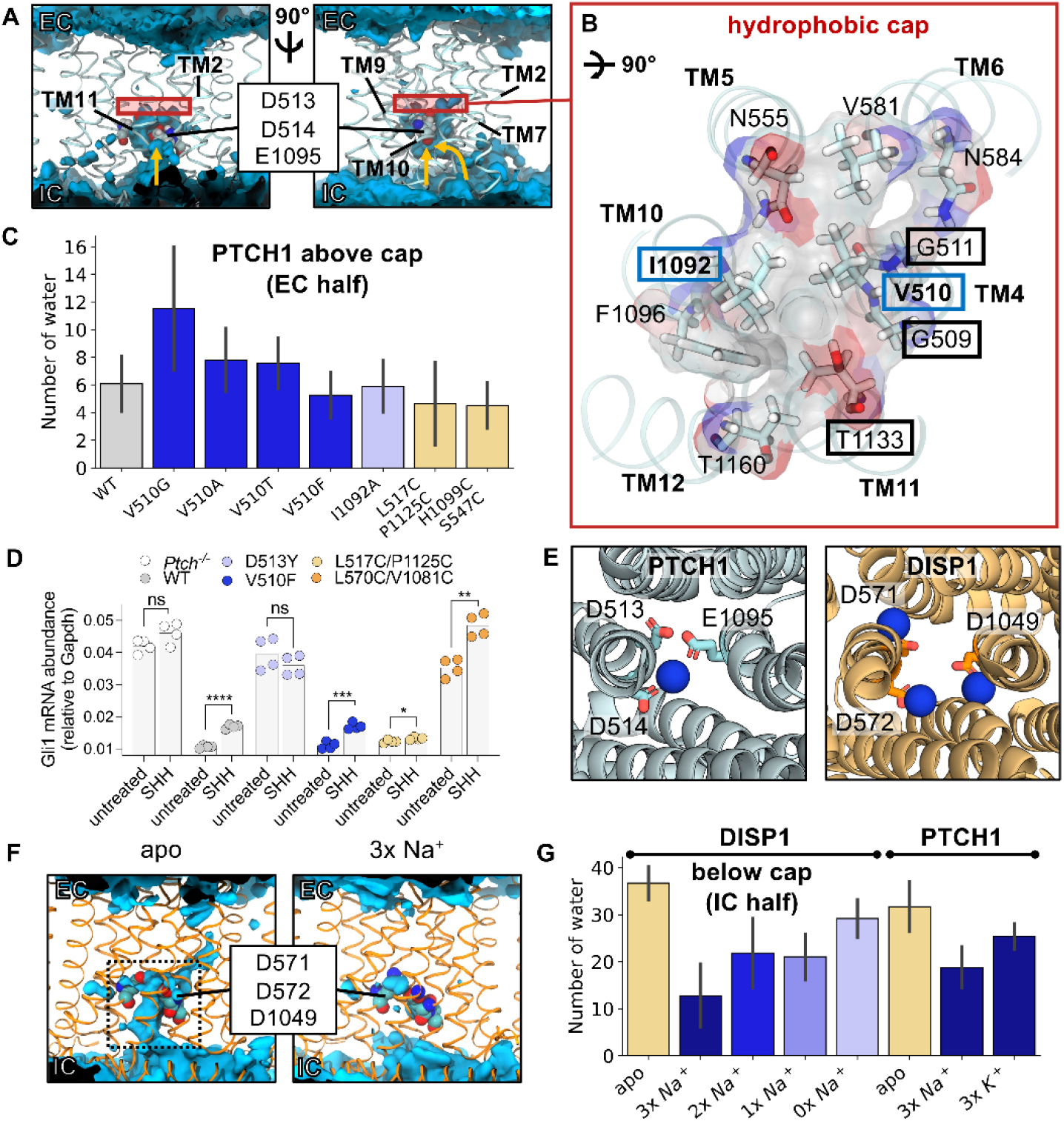
Identification of ‘inward-open’ and solvent ‘occluded’ PTCH1/DISP1 conformations. **A)** Time averaged water density (blue isosurface) across a 100 ns simulation of PTCH1 (PDB: 6DMY^5^) initiated with Na^+^ bound at Site 1. PTCH1 TMD is shown in ribbon representation and anionic triad residues are shown as spheres. Yellow arrows indicate paths for water entry into the TMD. **B)** Residues comprising the hydrophobic cap (red box in **A**) shown in stick and surface representation, as viewed from the extracellular (EC) face. V510 and I1092 forming part of the conserved GXXXDD and GXXX(E/D) motifs on TM4 and TM10 are boxed in blue. Residues mutated in disease phenotypes are boxed in black. **C)** Mean number of waters per frame within the EC half of the PTCH1 TMD across the final 10 ns of 3 x 50 ns simulations of WT PTCH1 or PTCH1 mutants (see Extended Methods). **D)** HH signalling strength is determined by measuring endogenous *Gli1* mRNA abundances (normalized to the control *Gapdh)* in response to SHH ligands (200 nM, 20 hours) in *Ptch1^-/-^* cells stably expressing the indicated variants. Statistical significance is determined by a Student’s t-test with a Welch’s correction. Exact *P* values for comparisons: *Ptch1^-/-^* untreated vs. SHH = 0.085, WT untreated vs. SHH < 0.0001, D513Y untreated vs. SHH = 0.2712, V510F untreated vs. SHH = 0.0002, L517C/P1125C untreated vs. SHH = 0.0255, and L570C/V1081C untreated vs. SHH = 0.0014. not significant (ns), \**P* > 0.05, \**P* ≤ 0.05, \*\**P* ≤ 0.01, \*\*\**P* ≤ 0.001, and *\*\*\*\*p* ≤ 0.0001. **E)** Comparison of cation binding sites within PTCH1 and DISP1 (PDB: 7RPH^44^) TMDs. Anionic triad residues are shown as sticks. **F)** Time averaged water density profiles across 100 ns simulations of DISP1 in apo or ‘3x Na^+^’ bound conformations. **G)** Mean number of waters per frame within the intracellular (IC) half of the DISP1 TMD across the final 10 ns of 3 x 100 ns simulations of DISP1 in apo and ‘3x Na^+^’ bound states, or in ‘2x Na^+^’, ‘1x Na^+^’ and ‘0x Na^+^’ bound states generated by sequential ion removal from the end of the previous Na^+^ bound state or PTCH1 in apo, ‘3x Na^+^’ and ‘3x K^+^’ bound states (see Extended Methods). **C** and **G** report the mean and standard deviation between repeats.

### Identification and mutation of the hydrophobic cap

Trajectory visualisation suggested water permeation into the extracellular region of the TMD may be prevented by several hydrophobic residues situated directly above the anionic triad (Fig. 5B). Notably two residues involved in formation of the hydrophobic cap, V510 and I1092, are each flanked by two glycine residues which could facilitate sidechain swivel to enable solvent permeation. These residues also comprise the conserved GXXXDD and GXXX(D/E) motifs on TM4 and TM10. To probe this hypothesis, we performed atomistic simulations of WT PTCH1 and PTCH1 TMD mutants without Na^+^ initially bound at Site 1. Residues constituting the hydrophobic cap (V510, I1092) were selected for *in silico* mutation in an attempt to promote solvent permeation through the TMD. In 14/18 (78 %) of trajectories Na^+^ entered the TMD and bound at Site 1, often within the first 20 ns of the simulations (Supplementary Fig. 9). We analysed the number of waters within the extracellular TMD half of a cylinder surrounding residues V510 and I1092 within the last 10 ns of each simulation (Fig. 5C). Compared to WT PTCH1 (waters: 6 ± 2) the average number of water molecules per frame within the extracellular half of the TMD was increased when V510 was mutated to glycine (waters: 12 ± 5) but not when mutated to a smaller hydrophobic (V510A, waters: 8 ± 2) or polar (V510T, waters: 8 ± 1) residue. The I1092A (waters: 6 ± 1) mutation also had no effect on solvation of the extracellular half of the TMD, nor did attempts to disrupt the packing of cap residues (V510F, waters: 5 ± 1). To functionally test the role of the V510 residue in forming a hydrophobic cap, we tested the V510F mutation in HH signalling assays. In cells lacking *Ptch1* (*Ptch1^-/-^*), high levels of unrestrained HH signalling are observed (Fig. 5D) compared to repressed SMO activity in cells expressing WT hPTCH1. Mutation of D513Y, proposed to coordinate Na^+^/K^+^ ion binding at Site 1 in PTCH1, was not functional to inhibit HH signalling. Next, we tested the activity of V510F, and found that it is functional to block HH signalling, and it demonstrates responsiveness to SHH ligand addition. This *in vivo* mutational data is consistent with the *in silico* data showing that water permeation was unaltered by the V510F mutation. We also expressed the V510G mutant in *Ptch1^-/-^* cells, predicted to have increased water permeation into PTCH1, but this mutant displayed lower expression by Western blot analysis (Supplementary Fig. 8). This may suggest that increased solvent permeability in PTCH1 reduces its stability.

Overall, our *in silico* and *in vivo* data suggest a model for solvent permeation governed purely by a simple hydrophobic restriction at one position on TM4/TM10 (as occurs in e.g. ion channels) is overly simplistic. This suggests ion coupling likely involves more complex concerted TMD motions.

### Breathing-like motions of the PTCH1 TMD mediate ion binding and PTCH1 function

We next tested whether mutations at the intracellular TMD face of PTCH1 could block solvent permeation. We tested the effect of introducing disulphide bonds at residues near the opening of the hydrophilic cavity to sterically prevent water entry. First, we cross-linked the intracellular TMD face on TM4-TM11 (L517C-P1125C, waters: 4 ± 3) or TM5-TM10 (H1099C-S547C, waters: 4 ± 1), which marginally decreased water permeation and, notably, prevented Na^+^ entry to TMD Site 1 in all but one trajectory (Fig. 5C, Supplementary Fig. 9). These cross-linked mutants support the proposal that breathing-like motions of the intracellular TMD regions, restricted in the cross-linked mutants, may regulate ion entry within PTCH1.

To further dissect the role of TMD motions in PTCH1 function we performed *in vivo* assays of PTCH1 mutants (Fig. 5D). Using the system described above, we expressed L517C-P1125C in *Ptch^-/-^* cells, and found that it was functional to inhibit SMO activity. Interestingly, this mutant was not responsive to SHH ligands, suggesting that conformational flexibility at the intracellular side of PTCH1 is required for its inactivation by SHH. Alternatively, the disulphide bond expected to form between the two residues may not occur when PTCH1 is in an active conformation with the open intracellular cavity, but may form in a conformational intermediate induced by SHH binding. Next, we tested a disulphide mutant on the extracellular side of the TMD, L570C/V1081C. Surprisingly, this mutant was not functional, despite being expressed at high levels, but did exhibit SHH responsiveness (Fig. 5D, Supplementary Fig. 8). This suggests that conformational flexibility in the extracellular face of the TMD are required for PTCH1 function. This type of movement may be required for ion movement from outside the cell towards the pocket or for release of ions within the pocket to the extracellular aqueous environment. Future work is needed to distinguish between these models.

### Insights from DISP1: Na^+^ binding induces transition to an occluded conformation

DISP1 is an RND transporter which catalyses the Na^+^ coupled export of SHH from the plasma membrane into the extracellular space (upstream of SHH engagement with PTCH1)^21,44,63^. The DISP1 TMD shares homology with PTCH1, and a recent DISP1 structure revealed three Na^+^ ions bound to the anionic triad within the TMD (Fig. 5E)^44^. We performed atomistic simulations of DISP1 in ‘3x Na^+^’ and apo states to assess whether a) three Na^+^ ions could be accommodated within this small, charge-dense region and b) whether DISP1 could inform our understanding of PTCH1 function. Analysis of the time-averaged water density within apo DISP1 revealed a solvated intracellular-open cavity, similar to PTCH1 (Fig. 5F). Surprisingly this intracellular cavity was absent in the ’3x Na^+^’ state and all three Na^+^ ions were stably coordinated within the TMD (RMSD: 0.16 ± 0.01 nm per Na^+^), validating the interpretation of cryo-EM densities^44^. We reasoned the 3 x Na^+^ ions may stabilise a solvent occluded state by interlocking neighbouring TM helices. To test this, TMD Na^+^ ions were sequentially removed from the end snapshot of simulations to generate successive ‘2x Na^+^’, ‘1x Na^+^’ and ‘0x Na^+^’ states. Cation removal increased the average number of waters per frame within the intracellular region of the TMD, indicating all three Na^+^ ions contribute to TMD stabilisation and restriction of water entry (Fig. 5G).

We calculated minimum distances between the intracellular portions of TM helices in DISP1. Compared to extracellular TM distances (which remined within ~0.1 nm regardless of Na^+^ bound state, Supplementary Fig. 10A), distances between intracellular TM portions were constantly increased in the apo state by ~0.4 nm relative to the ‘3x Na^+^’ state (Fig. 6). Sequential removal of ions increased intracellular TM distances towards those of the apo state. Thus, 3 x Na^+^ ion binding within the TMD of DISP1 facilitates transition from an ‘inward-open’ to solvent ‘occluded’ state by restricting outward-displacement of the intracellular regions of TM helices (Fig. 6D).

**Figure 6:**
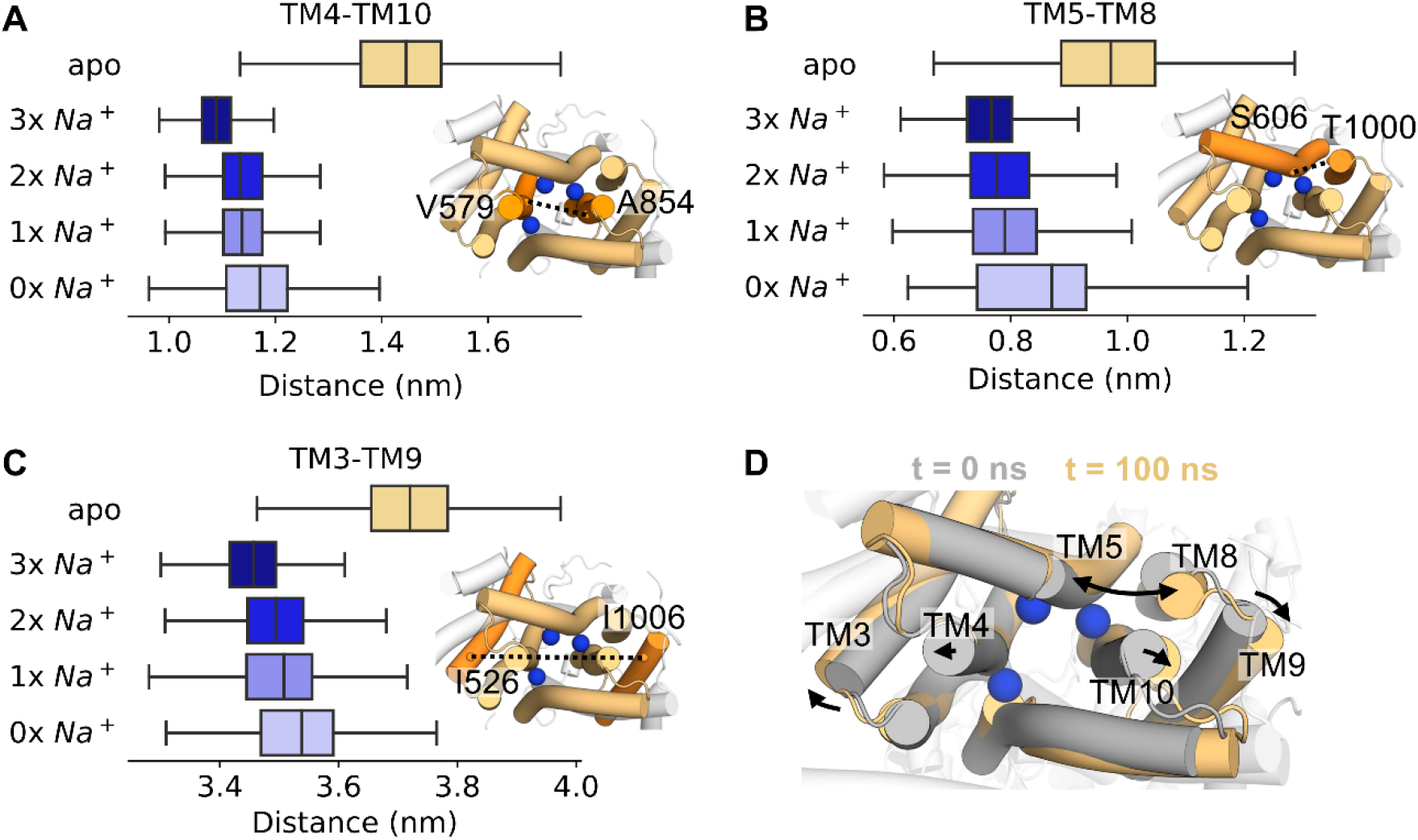
Na^+^ interdigitates the DISP1 intracellular TMD helices. **A-C)** Minimum distances between the intracellular portions of DISP1 helices across 3 x 100 ns simulations of DISP1 in apo, ‘3x Na^+^’, ‘2x Na^+^, ‘1x Na^+^’ and ‘0x Na^+^’ bound states. Distances were calculated between the Cα atoms of labelled residues on **A)** TM4-TM10, **B)** TM5-TM8 and **C)** TM3-TM9. Snapshots of the DISP1 TMD from the intracellular face are shown, with black lines marking the distance between helices (orange). **D)** Expansion of the intracellular TMD helices as indicated by comparison of the start (grey) and end (yellow) snapshots from a simulation of apo DISP1. The position of Na^+^ ions (blue spheres) are overlayed for reference.

Propelled by these findings we sought to assess whether cation mediated TMD interdigitation also occurs in PTCH1. We simulated PTCH1 with either ‘3x Na^+^’ or ‘3x K^+^’ ions bound at equivalent positions to in the DISP1 structure^44^. Na^+^/K^+^ ions remained bound albeit with a higher RMSD (0.33 ± 0.03 nm per Na^+^, 0.43 ± 0.05 nm per K^+^) than observed for DISP1 Na^+^ ions which may indicate further sidechain rearrangements are required for optimal coordination. For example, exchange with, or movement towards Site 2 was occasionally observed for the Na^+^/K^+^ ion coordinated by D513 (equivalent to DISP1 D571) (Supplementary Fig. 9). As in DISP1, the presence of 3 x Na^+^/K^+^ ions reduced water entry into the PTCH1 TMD cavity compared to when apo (Fig. 5G) and prevented TMD opening between ‘occluded’ (‘3x Na^+^/K^+^’) and ‘inward-open’ (apo or ‘1x Na^+^’ bound) conformations (Supplementary Fig. 10B-C). This corroborates our findings from the cross-linked *in silico* and *in vivo* PTCH1 mutants that subtle breathing-like motions play a role in solvent permeation through the TMD, and may also explain why ordered, trapped water molecules could be resolved with the DISP1 TMD^44^ but not in current PTCH1 structures.

## Discussion

Several models have been proposed for PTCH1 mediated SMO inhibition (Fig. 7A), extending from biochemical observations suggesting PTCH1 reduces accessible cholesterol levels in the cilia, and thus lowers SMO activation levels^1,14^. Models include extraction of cholesterol from the SMO cysteine-rich domain and import through PTCH1 to a sequestered cholesterol pool (such as in complex with sphingomyelin) or intracellular acceptor (model-1), export of accessible cholesterol through PTCH1 (model-2) or re-partitioning of accessible cholesterol via an ECD-dependant mechanism (model-3)^14,52^.

**Figure 7:**
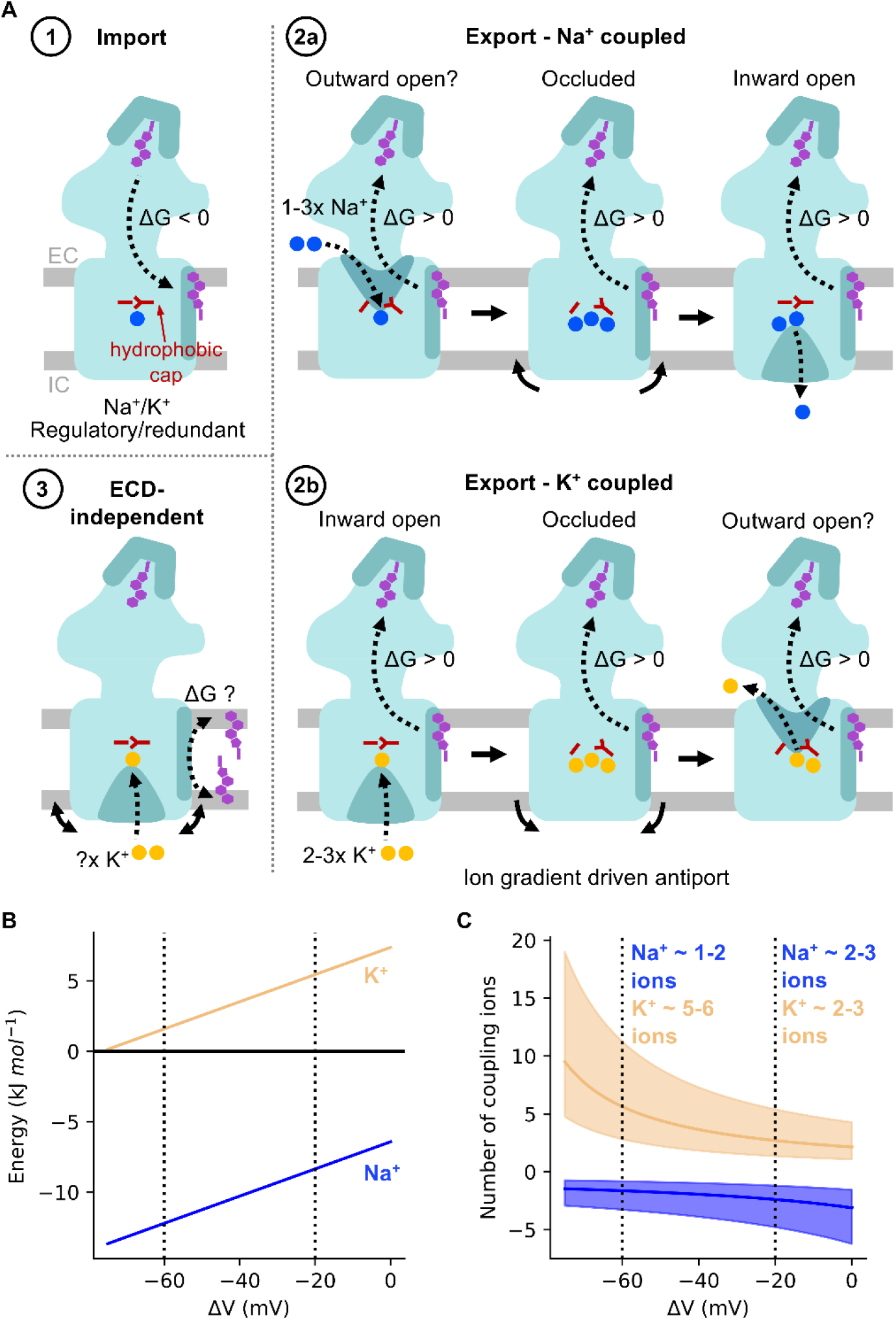
Energetics and ion coupling of PTCH1 transport models. **A)** Proposed models for PTCH1 function (coloured as in Fig. 1). Model-1: PTCH1 imports cholesterol extracted from SMO to a membrane sequestered pool or intracellular donor (energetically favourable). Model-2: Accessible cholesterol export by PTCH1 mediated by coupling to 1-3 Na^+^ ions (blue, model-2a) or 2-3 K^+^ ions (yellow, model-2b). Transition between ‘inward-open’ and ‘occluded’ states accompanied by breathing-like motions of intracellular helical segments (solid arrows) and alleviation of the hydrophobic cap (red). Model-3: PTCH1 re-partitions accessible cholesterol to an intramembrane sequestered pool (e.g. partnered with sphingomyelin) or intracellular acceptor. **B)** Free energy stored across the Na^+^ (blue) and K^+^ (yellow) membrane gradients as a function of membrane potential (ΔV) (Equation 2). **C)** Predicted number of Na^+^/K^+^ coupling ions required per cholesterol exported by PTCH1 (defined as ‘cholesterol export free energy’/’free energy across cation potential’ from **B**) *vs* membrane potential (ΔV). A cholesterol export free energy of +20 kJ mol^-1^ is indicated by a solid line, with export energies ranging between +10 to +40 kJ mol^-1^ indicated in transparent. Standard cellular ion concentrations ([Na^+^]_in_: 12 mM, [Na^+^]_out_: 145 mM, [K^+^]_in_: 150 mM, [K^+^]_out_: 4 mM) are assumed.

Using two independent free energy methods we predict cholesterol import (model-1) to be energetically favourable whereas movement in the export direction (model-2) would require coupling to an energy source. Model-1 is not consistent with the requirement of cations for PTCH1 function^21,22,52^ or with anionic triad mutants which render PTCH1 less functional and are associated with severe cooccurrences of Gorlin’s syndrome^5,51,64–66^. There are also no known extracellular cholesterol shuttles between SMO and PTCH1 which would presumably be necessary given their distinct membrane localisations^20,67^. The PTCH1 homologue NPC1^68–72^ can however, spontaneously transport cholesterol in the import direction^73^, which can be rationalised by our free energy calculations.

Our atomistic simulations suggest cholesterol export could be made favourable by coupling to translocation of Na^+^/K^+^ through the TMD (Fig. 4, Supplementary Fig. 3, 9). Biochemically, both Na^+22^ and K^+21,52^ have been implicated in PTCH1 function however assay interpretation is complicated by use of downstream signalling readouts for PTCH1 activity. K^+^ involvement is also perplexing given there are no known eukaryotic K^+^-coupled transporters and the K^+^ equilibrium potential approaches the predicted plasma membrane potential. Accordingly, our FEP calculations predict similar binding affinities for K^+^ and Na^+^ at Site 1 (Fig. 4E-F). To estimate the cholesterol:cation transport stoichiometry we calculated the free energy stored in the Na^+^/K^+^ gradient as a function of membrane potential (Fig. 7B, Equation 2). This was used to obtain the number of export-coupled cations, assuming a generous cholesterol export free energy of ΔG_1_= +10 to +40 kJ mol^-1^ and standard cellular ion concentrations ([Na^+^]_in_: 12 mM, [Na^+^]_out_: 145 mM, [K^+^]_in_: 150 mM, [K^+^]_out_: 4 mM) (Fig. 7C).

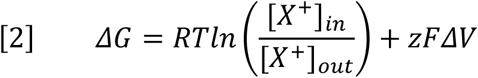

Thus, we predict 1-3 Na^+^ (model-2a) or 2-3 K^+^ (model-2b) ions to be required per cholesterol transported, although K^+^ driven transport could only occur if the cilia membrane potential was less electronegative than expected (approaching −20 mV) (Fig. 7C). There is some suggestion this may be the case^74^ however it is unclear how this would be established given the cilia is contiguous with the plasma membrane.

While a PTCH1 cholesterol export model is supported by ECD localised sterol densities in cryo-EM structures^5,8–10^, it is difficult to reconcile with one long-standing observation; PTCH1 ECD2 cleavage renders PTCH1 constitutively active^51,57^. A recently proposed ECD-independent mechanism (model-3) addresses this by suggesting PTCH1 may promote re-partitioning of accessible cholesterol to an intracellular acceptor or sphingomyelin sequestered pool^52^. Hence the ECD may simply be a socket for SHH engagement independent of PTCH1 function. Our free energy calculations support a role for the SBD as an energetic switch between ‘SHH-cholesterol’ and ‘free cholesterol’ engaged conformations (Fig. 2D). The free energy of interleaflet cholesterol flippase activity would presumably be much lower (given this occurs spontaneously in membranes^75^) and thus may be more amenable to reduced free energy stored in the K^+^ potential at ~-60 mV. Thus, we turn the export argument on its head by asking whether the incompatibility of the K^+^ and cholesterol export free energies (at ~-60 mV, Fig. 7B-C) can be used as further evidence against an ECD-dependant PTCH1 function? This is consistent with our data for the ECD salt-bridge (P115R/N802E) retaining its ability to block HH signalling while losing SHH responsiveness (Fig. 2E).

Finally, we complemented our PTCH1 analyses with simulations of DISP1 in distinct Na^+^ bound states^44^ (Fig. 5-6). In the ‘3x Na^+^’ state solvent is prevented from entering DISP1 by Na^+^ mediated interdigitation of the intracellular portions of TM helices. Ion removal promoted transition from a solvent ‘occluded’ to ‘inward-open’ conformation by concerted outward movement of intracellular helical segments (Fig. 6). This effect was also observed when PTCH1 was simulated in the presence of 3 x Na^+^/K^+^ ions (Fig. 5, Supplementary Fig. 10). Thus, we provide the first molecular description of the conformational transition between distinct ion-coupled transport states of DISP1/PTCH1. More work is needed to establish the events leading to putative ‘outward-open’ conformations and/or cross-membrane ion translocation.

In conclusion, we provide energetic contextualisation to the directionality and stoichiometry of PTCH1 transport models, and elucidate conformational transitions pertinent to other RND protein mechanisms.

## Acknowledgments

The authors thank Dr Owen Vickery for guidance with PMF calculations and use of the *pmf.py* tool and Dr Wanling Song for use of the PyLipID toolkit. T.B.A., R.A.C. and M.S.P.S are supported by Wellcome (102164/Z/13/Z and 208361/Z/17/Z). M.S.P.S.’s research is additionally supported by the BBSRC (BB/R00126X/1) and PRACE (Partnership for Advanced Computing in Europe, 2016163984). C.S. is supported by Cancer Research UK (C20724/A26752) and the BBSRC (BB/T01508X/1). R.R. is supported by the National Institutes of Health (GM118082 and GM106078). M.K. is in receipt of a pre-doctoral fellowship from the National Science foundation. L.V.V. is supported by a Cancer Research UK studentship (C20724/A27829). This project made use of ARCHER supercomputing time under the UK High-End Computing Consortium for Biomolecular Simulation (HECBioSim; EPSRC EP/R029407/1).

## Author contributions

T.B.A. and R.A.C. performed and analysed simulations. L.V. and C.S. cloned and expressed PTCH1 mutants. M.K and R.R. performed PTCH1 functional assays. M.S.P.S. and C.S. conceptualised the project. T.B.A., R.A.C. and M.S.P.S. wrote the paper with input from all authors.

