## Supplementary for "The Energetics and Ion Coupling of Cholesterol Transport Through Patched1"

#### Supplementary Information

##### Extended Methods:

Setup of coarse-grained potential of mean force calculations

The following parameters were used in all CG simulations unless otherwise stated. The MARTINI 2.2 forcefield was used to describe all components[1,2]. Protein coarse-graining was achieved using the *martinize.py* script with the ElNeDyn elastic network, a spring force constant of 500 kJ $mol^{-1}$ $nm^{-2}$ and a cut-off of 0.9 nm [3]. Membranes were built using *insane.py*[4]. All systems were solvated using MARTINI water and approximately 0.15 M NaCl. A velocity (V-)rescaling thermostat[5] was used to maintain temperature at 310 K ($\tau_t$=1.0 ps). The Parrinello-Rahman barostat[6] was used to maintain pressure at 1 bar ($\tau_p$=12.0 ps) with a compressibility of $3x10^{-4}$ $bar^{-1}$. The timestep was 20 fs.

*PMF-1: between the SSD and SBD and PMF-2: between the SSD and bulk membrane*

The SSD of PTCH1-molA and PTCH1-molB (PDB: 6RVD[7]) were aligned to the x axis to assist generation of the reaction coordinate and the surrounding membrane and solvent expanded to a box size of 20 x 20 x 19 $nm^3$ using gmx genconf and VMD[8]. Two round of steepest decent energy minimisation were performed to relax the equilibrated bilayer around PTCH1. For PMF-1a the 'SHH-cholesterol' or 'free cholesterol' molecules were positioned in PTCH1-molA and PTCH1-molB ECDs respectively according to the cryo-EM densities[7]. PMF-1b was initiated with cholesterol bound at the base of the ECD, continuing from the final window in PMF-1a from PTCH1-molB. Since the 'free cholesterol' bound within PTCH1-molB was positioned with the ROH bead orientated towards the bilayer, a bash script was used to invert the cholesterol orientation in frames generated from the steered MD simulations to generate the PTCH1 'SHH-cholesterol' conformations. This was due to difficulties obtaining a reasonable steered MD simulation initiated from the final window of PMF-1a from PTCH1-molA and to standardise the reaction coordinate. For PMF-2 cholesterol was not present in the PTCH1 ECD. Instead, snapshots were selected with cholesterol bound to the SSD with either the ROH bead (equivalent to the $\beta_3$-OH group) pointing towards the extracellular leaflet headgroups ('OH-up') or towards the bilayer midplane ('OH-down').

*PMF-3: between the bulk membrane and solvent*

A 10 x 10 x 18 $nm^3$ POPC:CHOL (3:1) bilayer patch without protein was built using *insane.py*[4,9]. The bilayer was solvated using MARTINI water[1] and 0.15 M NaCl before two round of steepest decent energy minimisation. A cholesterol molecule at the centre of the bilayer was selected for use in PMF calculations.

*PMF-4: between the SBD and solvent*

The ECD of PTCH1-molA and PTCH1-molB (residues A119–D436 and R772–G1023) from the PTCH1-SHH (2:1) structure (PDB: 6RVD[7]) was extracted and coarse-grained[1]. The 'SHH-cholesterol' and 'free cholesterol' molecules were positioned in the SBD of PTCH1-molA and PTCH1-molB respectively accordioning to their positions in the cryo-EM density[7] (and identically to in PMF-1a). The PTCH1 ECDs were positioned at one end of a 9 x 9 x 18 $nm^3$ box with the SBD aligned to the z axis. The system was solvated with MARTINI water[1] and approximately 0.15 M NaCl followed by two rounds of steepest decent energy minimisation.

Execution and analysis of coarse-grained potential of mean force calculations

Steered MD simulations were used to generate 1D reaction coordinates for each path of the PTCH1-molA and PTCH1-molB free energy cycles. The 1D reaction coordinate was generated by application of a distance dependant pulling force along specified axes between the COM of the cholesterol and the following selections: PMF-1a (I1092 BB bead), PMF-1b (F800 BB bead), PMF-2 (P504 BB bead), PMF-3 (COM bilayer) and PMF-4 (S331 BB bead). The umbrella pulling force was 1000 kJ $mol^{-1}$ $nm^{-2}$ and pulling rates of 0.1 nm $ns^{-1}$ (PMF-1a, 2, 3, 4) or 10 nm $ns^{-1}$ (PMF-1b) were used. Position restraints of 1000 kJ $mol^{-1}$ $nm^{-2}$ were applied to the following backbone beads to prevent protein rotation: PMF-1a/b, (A1088, A1157), PMF-2 (A1088, A1157) and PMF-4 (A182, A239). Frames were taken with spacing 0.05 nm along the reaction coordinate and subjected to 1-3 µs of simulation. For each window an umbrella pulling force of 1000 kJ $mol^{-1}$ $nm^{-2}$ was used to confine the cholesterol position along the reaction coordinate. Further details of the number of windows, simulations times and PMF convergence are given in Supplementary Table 1 and Supplementary Fig. 2. PMF profile analysis was assisted using the *pmf.py* tool[10] and the weighted-histogram analysis method[11] implemented in GROMACS (bootstrapped 2000 times).

Setup for atomistic simulations of PTCH1

A PTCH1 structure with ion-like density within the TMD (PDB: 6DMY)[12] was used in atomistic simulations. SHH, ligands and metal ions were removed. A short linker was modelled between TM6 and TM7 using Modeller 9.20[13] to give an overall sequence of (TM6)DRR-LDIFCC//TKWTLSSFAE-KHY(TM7), in accordance with the linker used in CG simulations of PTCH1 (PDB: 6RVD[7]). A $Na^+$ ion was positioned according to the density in the centre of the TMD[12]. The H++ server and propKa[14,15] were used to predict the pKa of titratable groups (Supplementary Table 4), revealing at least two residues of the anionic triad to be protonated at pH 7 in the absence of bound $Na^+$, decreasing to one protonated residue when $Na^+$ was present. The central residue, E1095, was therefore protonated using the CHARMM-GUI PDB generator which was also used to rename atoms to be compatible with the CHARMM-36 forcefield and model disulphide bonds between C203-C226, C234-C327, and C296-C304[16,17]. PTCH1 was embedded in a symmetric POPC:CHOL (3:1) membrane generated using the CHARMM-GUI bilayer builder[18,19] and solvated using TIP4P water[20] and approximately 0.15 M NaCl. PTCH1 was subsequently energy minimised and equilibrated in 2 x 5 ns NVT and NPT steps with restraints applied to the PTCH1 backbone.

*Application of a membrane potential*

Atomistic simulations of the above setup were also run in the presence of a -100 mV or -200 mV membrane voltage ($\Delta V = E.L_z$, E=-7.1429 or –14.286 mV, $L_z$=14 nm) for 3 x 100 ns each. Voltage is reported as inside relative to outside and was achieved using the constant electric field method[21].

*In silico mutation screen*

Modeller 9.20 was used to induce single amino acid substitutions at a specified location and induce disulphide bond formation in the cross-linked mutants[13]. The following mutations were performed: V510G, V510A, V510T, V510F, I1092A, L517C-P1125C, H1099C-S547C. Atomistic simulations of WT PTCH1 and each PTCH1 mutant were performed as described above with the exception that $Na^+$ was not initially bound in the TMD and each system was simulated for 3 x 50 ns.

Setup for atomistic simulations of DISP1

The DISP1 structure with three $Na^+$ ions bound was used in simulations (PDB: 7RPH, 'R conformation')[22]. Detergent ligands were removed, and the missing loop modelled using Modeller9.20[13]. CHARMM-GUI was used to generate DISP1 parameters (separated into two chains to account for the Furin cleavage site) and the anionic triad residue D1049 was protonated for consistency with the PTCH1 simulations[16,17]. DISP1 was simulated in the '3x $Na^+$' bound and apo states for 3 x 100 ns each. The '2x $Na^+$' state was generated from the end snapshots of the '3x $Na^+$' state by removing the ion missing in an alternative 'T conformation' of DISP1 (PDB: 7RPI)[22] bound to two $Na^+$ ions. The '1x $Na^+$' and '0x $Na^+$' states were generated by an identical process of sequential ion removal from the end snapshots of previous simulations. The ion retained in the '1x $Na^+$' state was chosen based on the apo state simulations whereby an ion spontaneously bound to D572 in all replicates. Each subsequent ion state was also simulated for 3 x 100 ns.

Analysis

*Protein-lipid interactions*

The PyLipID analysis toolkit[23] was used to identify residues in the PTCH1 ECD which interacted with ECD bound cholesterol (https://github.com/wlsong/PyLipID). All windows in the PMF-1a profile or windows surrounding energetic bottlenecks in the PMF-1a profile were analysed collectively using PyLipID with a single 0.6 nm interaction cut-off. The top four residues with the highest occupancy i.e. those residues within 0.6 nm of cholesterol for the largest fraction of the window simulation times were reported for each bottleneck.

*Water within PTCH1 and DISP1*

MDAnalysis[24] was used to analyse water within PTCH1 TMD. For each timepoint in the trajectory the O atom of water within a cylinder (radius 1.3 nm in xy) centred around the Cα atoms of residues V510 and I1092 was considered to localise within the TMD. For analysis of water z coordinates with time the length of the cylinder was 4 nm in z whereas for average water densities in the mutant screen a 3 nm cylinder was used to reduce contributions from bulk water molecules. Water within the extracellular and intracellular halves of the TMD was defined as having z coordinate of the O atom localised within the cylinder and either above or below the z coordinate of the midpoint between the Cα atoms V510 and I1092. Only the final 10 ns of each 50 ns simulation of PTCH1 were used in comparison of TMD water density to allow water to equilibrate in the TMD. PyMol and VMD were used to visualise trajectories[8].

Analysis of water within the intracellular TMD half of DISP1 was performed identically to described above with the cylinder midpoint defined using the Cα atoms of I568 and L1046.

### Supplementary Tables

**Supplementary Table 1:** Summary of the number of windows, simulations time per window and free energy value obtained from PTCH1 CG PMF calculations. The number of windows correspond to those included in the *gmx wham* step.

| PMF | System | No. windows | Window length (μs) | Total simulation time (μs) | ΔG (kJ $mol^{-1}$) |
|---|---|---|---|---|---|
| 1a | PTCH1-molA 'SHH-cholesterol' | 92 | 3 | 276 | +20 ± 4 |
| | PTCH1-molB 'free cholesterol' | 99 | 2 | 198 | -15 ± 7 |
| 1b | PTCH1-molA 'SHH-cholesterol' | 98 | 2 | 196 | +48 ± 9 |
| 2 | SSD Cholesterol 'OH up' PTCH1-molA | 65 | 1 | 65 | +18 ± 3 |
| | SSD Cholesterol 'OH up' PTCH1-molB | 56 | 1 | 56 | +16± 4 |
| | SSD Cholesterol 'OH down' PTCH1-molB | 65 | 1 | 65 | +4 ± 3 |
| 3 | Membrane cholesterol | 76 | 1 | 76 | +88 ± 1 |
| 4 | PTCH1-molA ECD 'SHH-cholesterol' | 112 | 2 | 224 | -83 ± 3 |
| | PTCH1-molB ECD 'free cholesterol' | 101 | 2 | 202 | -90 ± 4 |

**Supplementary Table 2:** Summary of the replicates, lambda values and free energies obtained from CG ABFE calculations.

| System | Replicates | No. λ windows | Window length (μs) | Total simulation time (μs) | ΔG (kJ $mol^{-1}$) |
|---|---|---|---|---|---|
| PTCH1-molA ECD 'SHH-cholesterol' | 3 | 29 | 0.15 | 13.05 | +92 ± 3 |
| PTCH1-molB ECD 'free cholesterol' | 3 | 29 | 0.15 | 13.05 | +86 ± 1 |
| PTCH1-molA SSD Cholesterol 'OH up' | 3 | 29 | 0.15 | 13.05 | +105 ± 1 |
| PTCH1-molA ECD 'mid cholesterol' | 3 | 29 | 0.15 | 13.05 | +66 ± 1 |
| PTCH1-molB ECD 'mid cholesterol' | 3 | 29 | 0.15 | 13.05 | +61 ± 2 |

**Supplementary Table 3:** Summary of equilibrium atomistic simulations of PTCH1 and DISP1.

| System | PDB | Replicates x Time | Initial ligands/conditions |
|---|---|---|---|
| PTCH1 | 6DMY | 3 x 100 ns | $Na^+$ (Site 1) |
| PTCH1 | 6DMY | 3 x 50 ns | apo (in 0.15 M NaCl) |
| PTCH1 | 6DMY | 3 x 100 ns | apo (in 0.15 M KCl) |
| PTCH1 | 6DMY | 3 x 100 ns | $Na^+$ (Site 1), -100 mV membrane potential |
| PTCH1 | 6DMY | 3 x 100 ns | $Na^+$ (Site 1), -200 mV membrane potential |
| PTCH1-mutants | 6DMY | 3 x 50 ns x 7 mutants | apo (in 0.15 M NaCl) |
| PTCH1 | 6DMY | 3 x 100 ns | '3x $Na^+$' |
| PTCH1 | 6DMY | 3 x 100 ns | '3x $K^+$' |
| DISP1 | 7RPH | 3 x 100 ns | apo |
| DISP1 | 7RPH | 3 x 100 ns | '3x $Na^+$' |
| DISP1 | 7RPH | 3 x 100 ns | '2x $Na^+$' |
| DISP1 | 7RPH | 3 x 100 ns | '1x $Na^+$' |
| DISP1 | 7RPH | 3 x 100 ns | '0x $Na^+$' |

**Supplementary Table 4:** pKa values of the PTCH1 anionic triad residues as predicted using the propKa and H++ servers.

| Residue | Predicted pKa | | |
|---|---|---|---|
| | **H++** | **H++ ($Na^+$ bound)** | **PropKa** |
| D513 | >12.00 | >12.00 | 8.45 |
| D514 | 6.26 | <0.00 | 8.24 |
| E1095 | 11.29 | 0.90 | 10.95 |

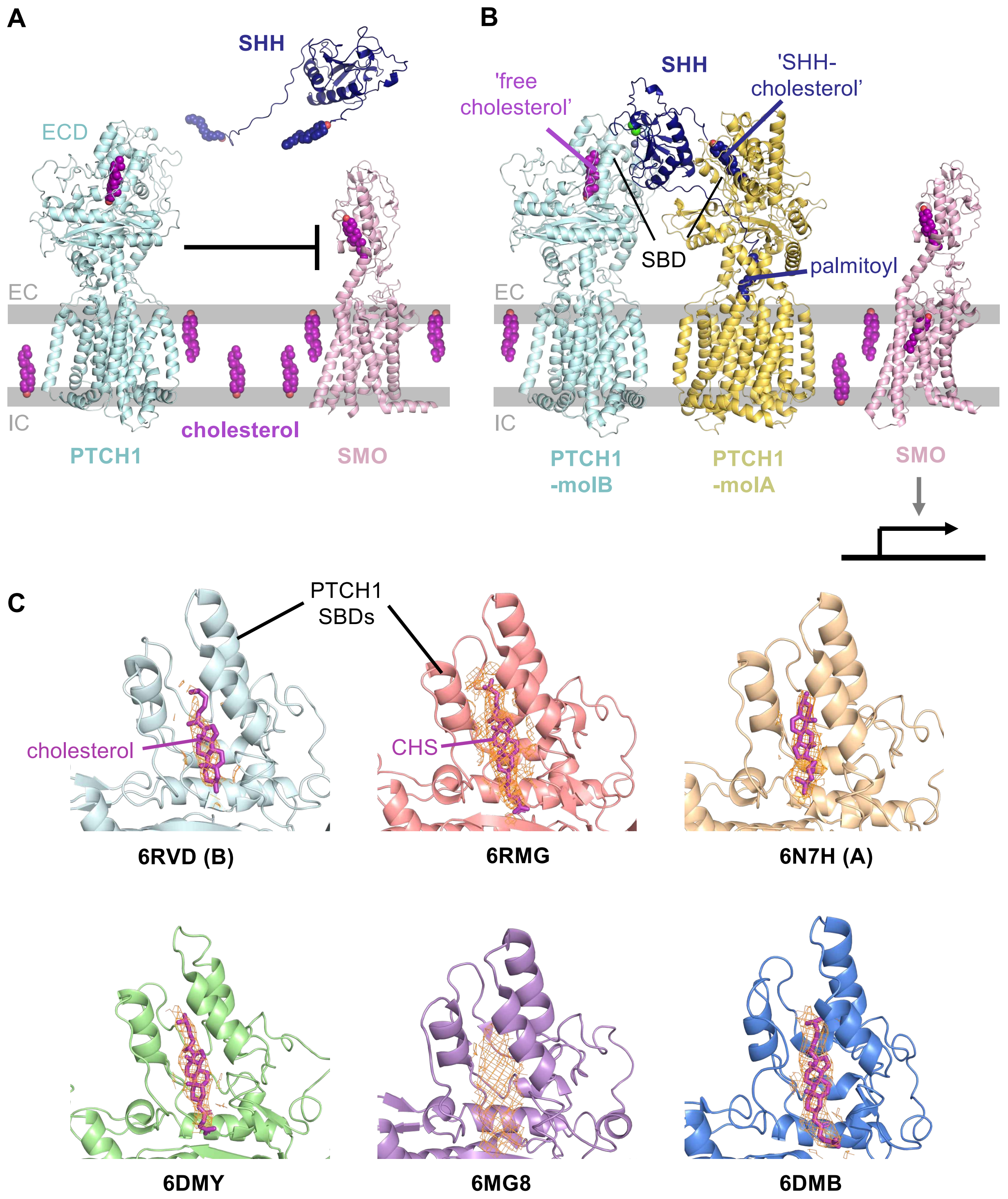


**Supplementary Figure 1: Patched1 and the vertebrate Hedgehog signalling pathway. A)** Patched1 (PTCH1, light blue) inhibits Smoothened (SMO, pink) in the absence of Sonic Hedgehog (SHH, dark blue). **B)** SHH binds to PTCH1 via a metal binding interface (PTCH1-molB) or insertion of terminal cholesteryl/palmitoyl attachments (PTCH1-molA, yellow), the later relieves SMO inhibition and initiates Hedgehog (HH) signalling. Proteins are shown in cartoon representation and cholesterol/palmitate are shown as spheres. Extracellular (EC) and intracellular (IC) leaflets are indicated. **C)** Cholesterol and cholesterol-hemisuccinate densities (orange) within the sterol binding domain (SBD) of existing PTCH1 structures. Modelled sterols are shown in purple.

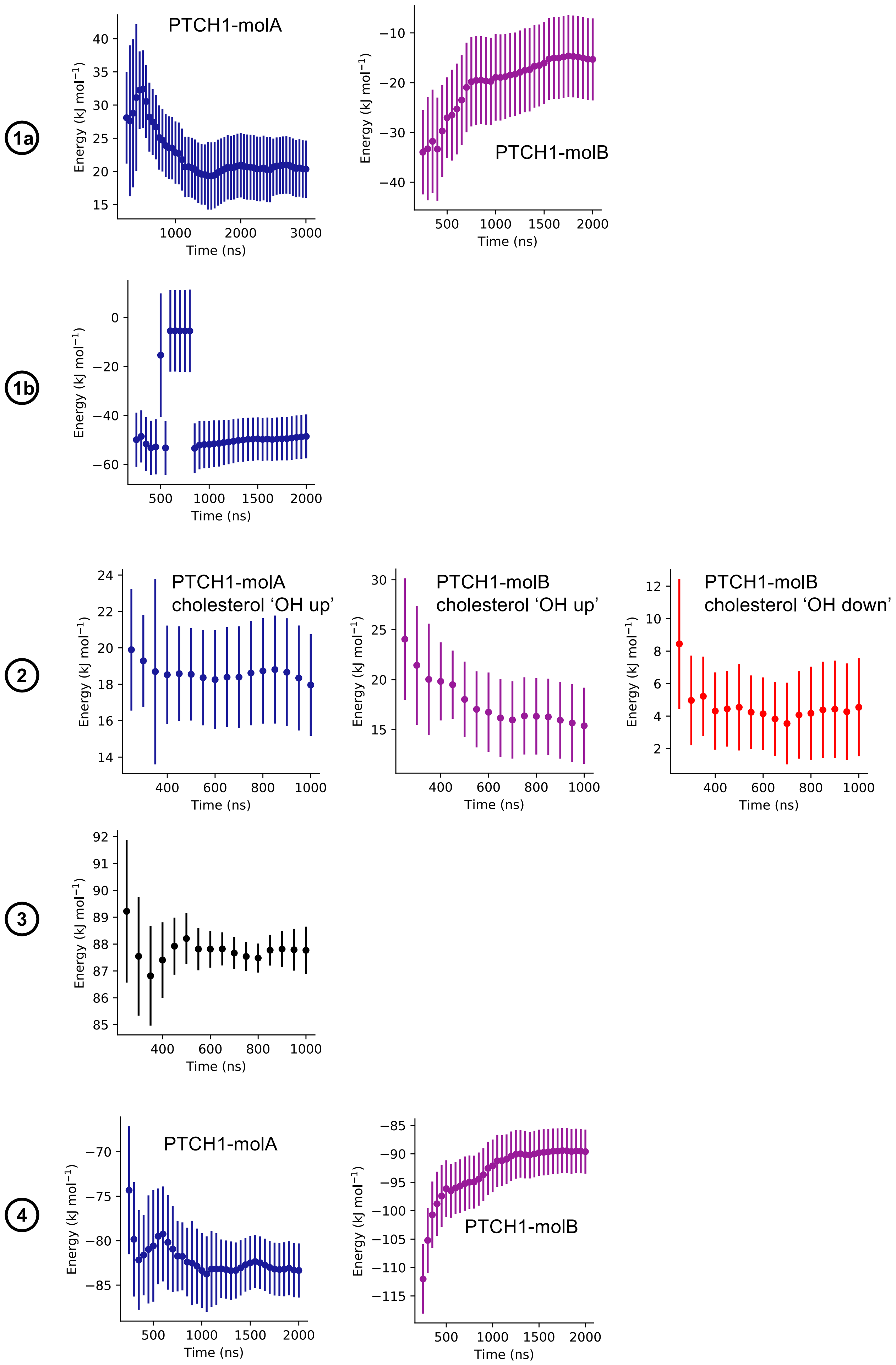
1a
PTCH1-molA
Energy (kJ mol⁻¹)
Time (ns)
PTCH1-molB
Energy (kJ mol⁻¹)
Time (ns)
1b
Energy (kJ mol⁻¹)
Time (ns)
2
PTCH1-molA
cholesterol ‘OH up’
Energy (kJ mol⁻¹)
Time (ns)
PTCH1-molB
cholesterol ‘OH up’
Energy (kJ mol⁻¹)
Time (ns)
PTCH1-molB
cholesterol ‘OH down’
Energy (kJ mol⁻¹)
Time (ns)
3
Energy (kJ mol⁻¹)
Time (ns)
4
PTCH1-molA
Energy (kJ mol⁻¹)
Time (ns)
PTCH1-molB
Energy (kJ mol⁻¹)
Time (ns)

**Supplementary Figure 2: Convergence of PMF calculations.**

Free energy from PMF calculations as a fraction of the per window simulation time analysed. Free energy values were calculated using gmx wham with 200 Bayesian Bootstraps, discarding the first 200 ns of each window as equilibration time. PMFs are numbered according to Fig. 1-2 and coloured according to ‘SHH-cholesterol’ (blue, bound to PTCH1-molA) and ‘free cholesterol’ (purple, bound to PTCH1-molB) orientations.

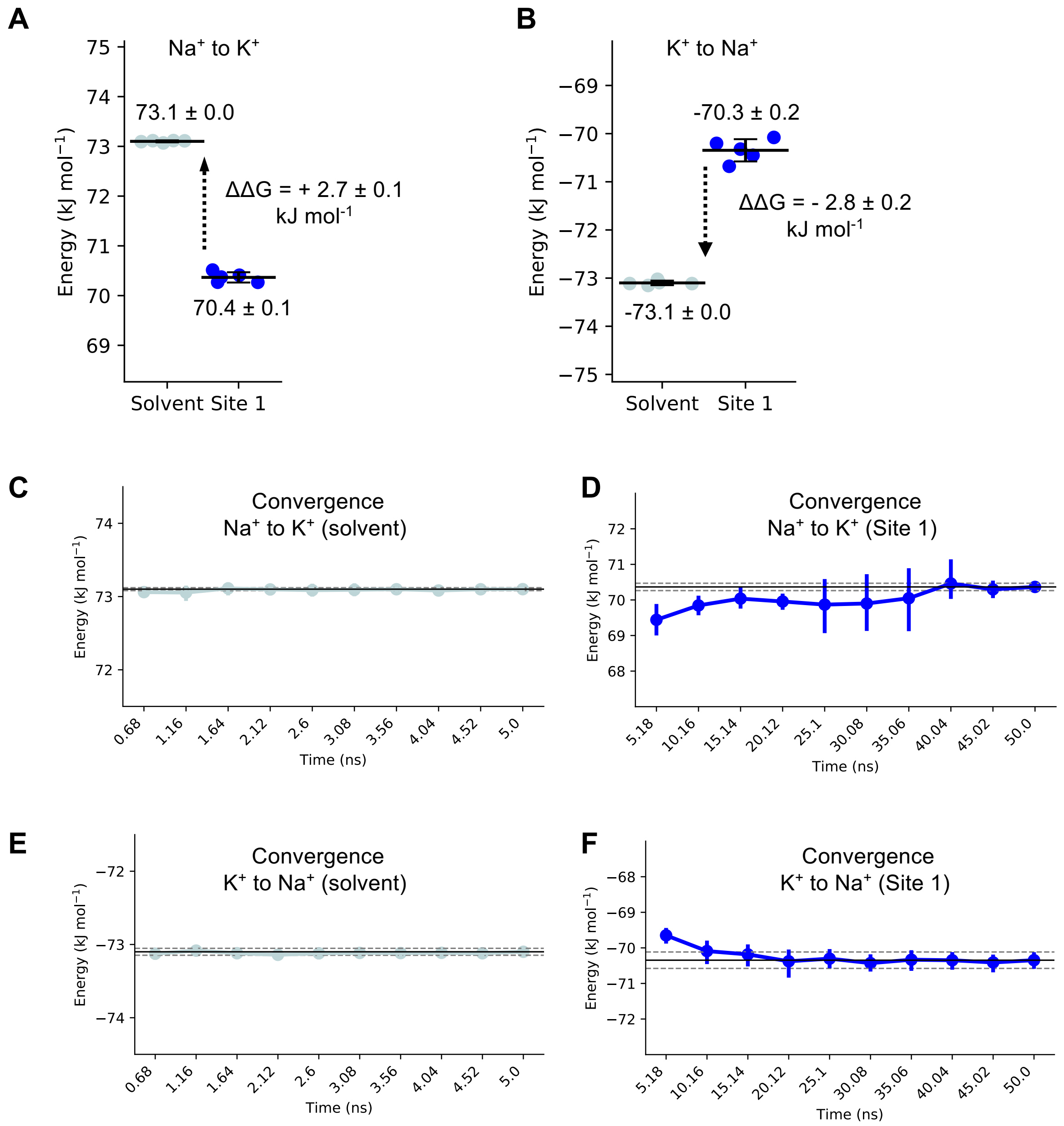


**Supplementary Figure 3: Convergence of FEP calculations.** Free energy perturbation of **A)** $Na^+$ to $K^+$ or **B)** $K^+$ to $Na^+$ when bound to PTCH1 Site 1 or in solvent. The forward and reverse perturbations are in agreement. **C-F)** Convergence of FEP calculations as a fraction of λ window length for **C)** $Na^+$ to $K^+$ in solvent, **D)** $Na^+$ to $K^+$ at PTCH1 Site 1 **E)** $K^+$ to $Na^+$ in solvent and **F)** $K^+$ to $Na^+$ at PTCH1 Site 1. Error bars indicate the minimum/maximum values (**A/B**) or standard deviation between 5 FEP replicates (**C-F**).

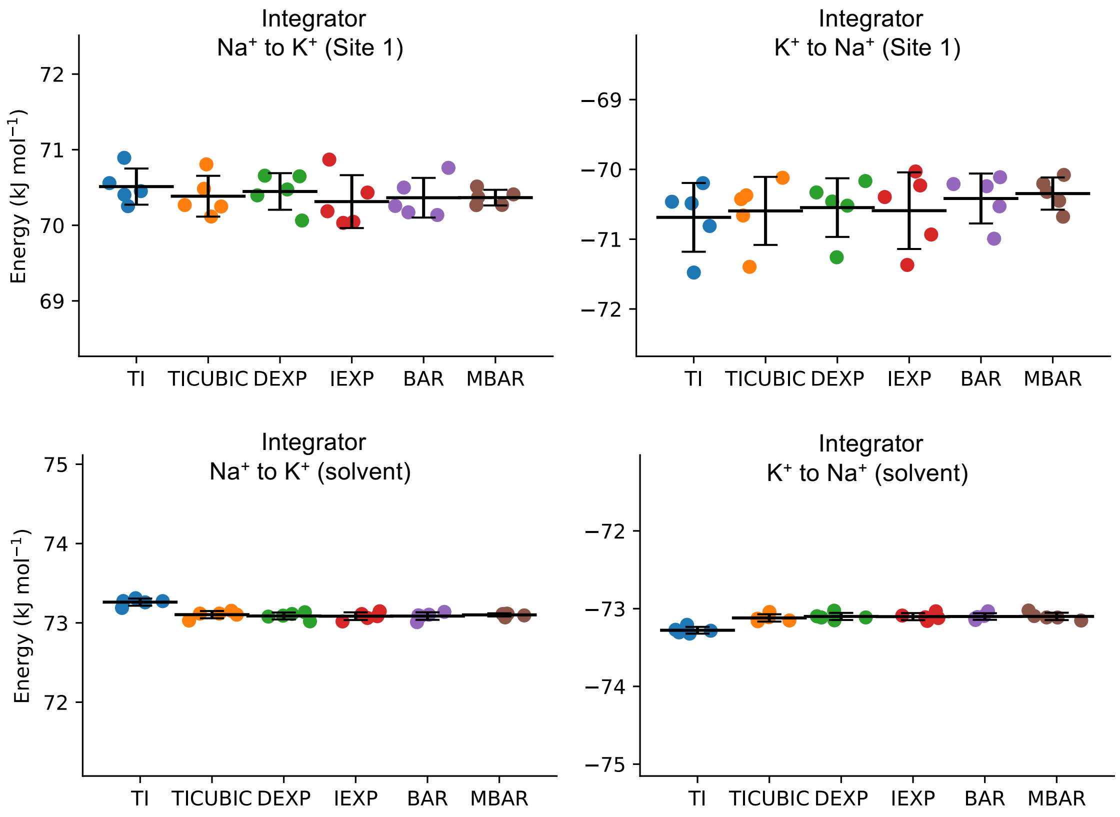


**Supplementary Figure 4: Choice of integrator in FEP calculations.** FEP values ion perturbations predicted using different integrators. See Supplementary Figure 3 legend for FEP details. The MBAR integrator was used to report FEP values.

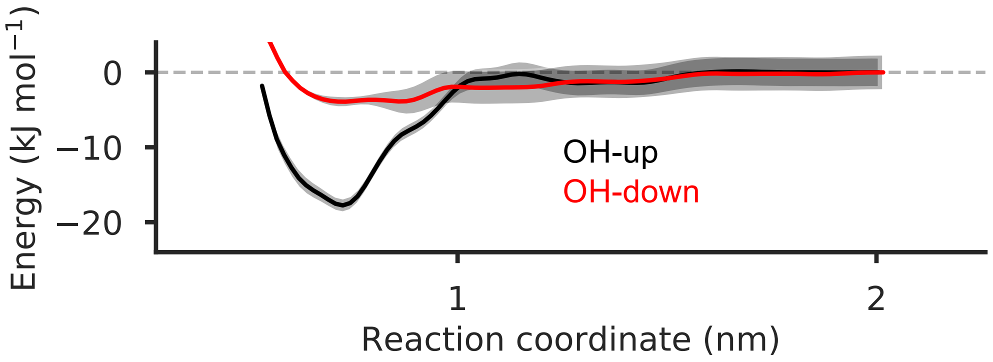


**Supplementary Figure 5: Cholesterol binds the PTCH1 SSD in the OH-up orientation.**

Coarse-grained (CG) potential of mean force (PMF) profiles for movement of cholesterol away from the PTCH1 sterol sensing domain (SSD) when bound with the ROH bead (equivalent to the β3-hydroxyl group) facing towards the lipid phosphate groups (OH-up, black) or towards the bilayer midplane (OH-down, red). Bayesian bootstrapping (2000 rounds) was used to estimate profile errors (grey).

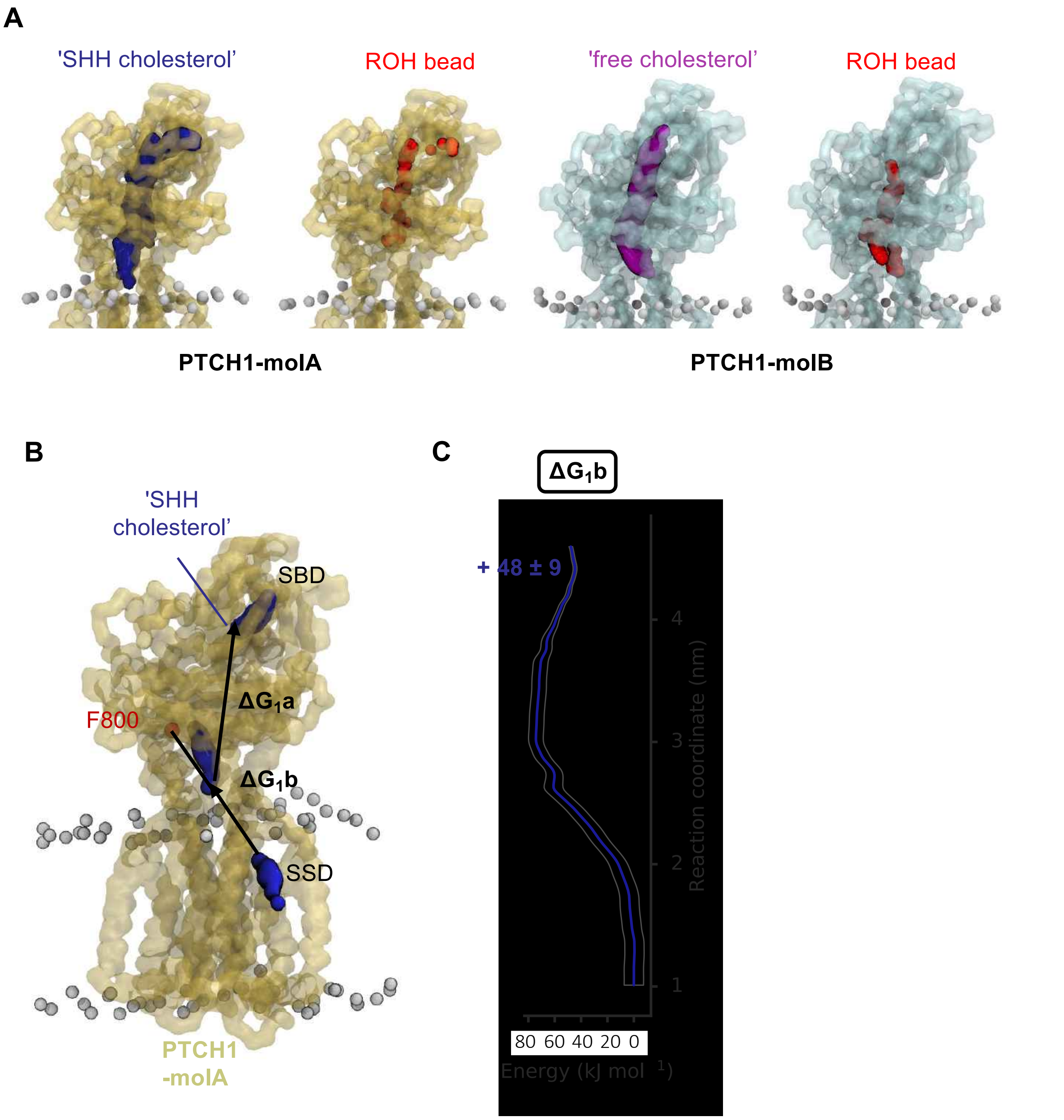


**Supplementary Figure 6: Cholesterol pathways through the PTCH1 ECD in PMF calculations.**

**A)** Coarse-grained (CG) representation of PTCH1-molA (yellow) and PTCH1-molB (blue) overlayed with the position of the 'SHH cholesterol' (dark blue) and 'free cholesterol' molecules, or their associated ROH beads (red) across all PMF-1a windows. **B)** Snapshots indicating movement of cholesterol between the sterol sensing domain (SSD) and the base of PTCH1 ECD in PMF-1b (residues used in the steered MD are labelled in red) in CG PTCH1-molA (yellow). **C)** PMF profile for cholesterol movement between the SSD and ECD base ($\Delta G_1b$). Bootstrapping errors (2000 rounds) are shown in grey.

A

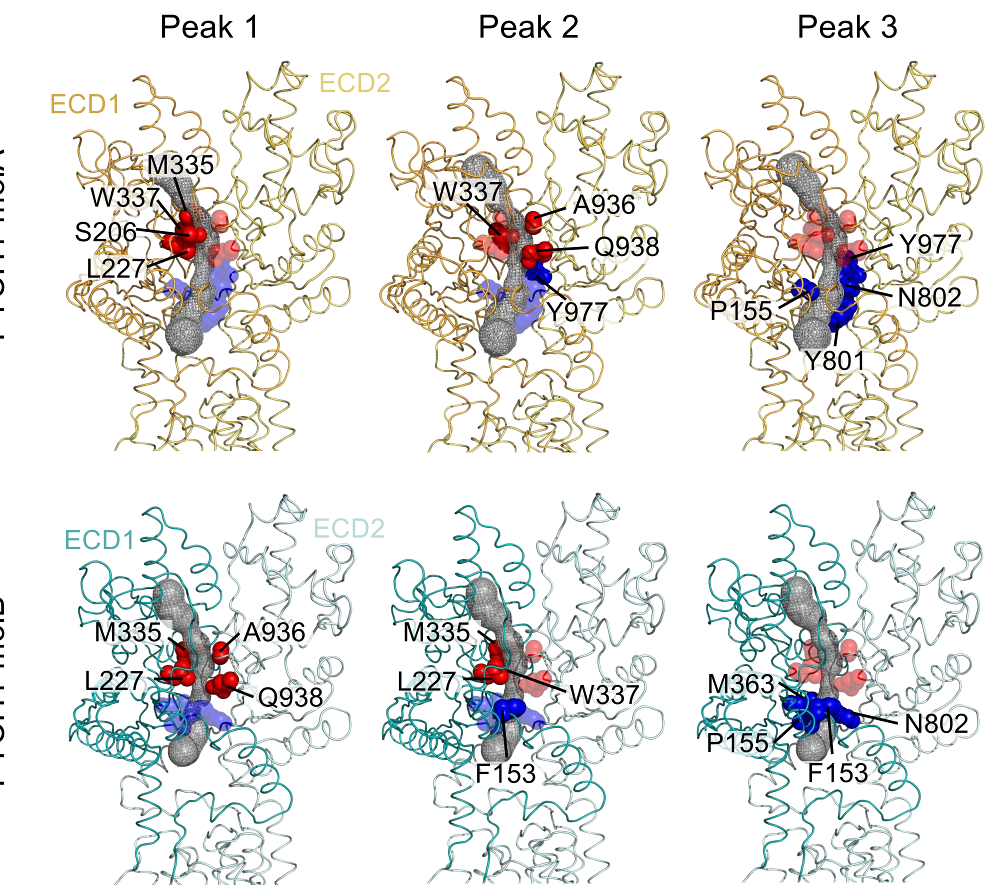
Peak 1
Peak 2
Peak 3
PTCH1-molA
ECD1
ECD2
M335
W337
S206
L227
W337
A936
Q938
Y977
Y977
N802
P155
Y801
PTCH1-molB
ECD1
ECD2
M335
A936
L227
Q938
M335
L227
W337
F153
M363
N802
P155
F153

B

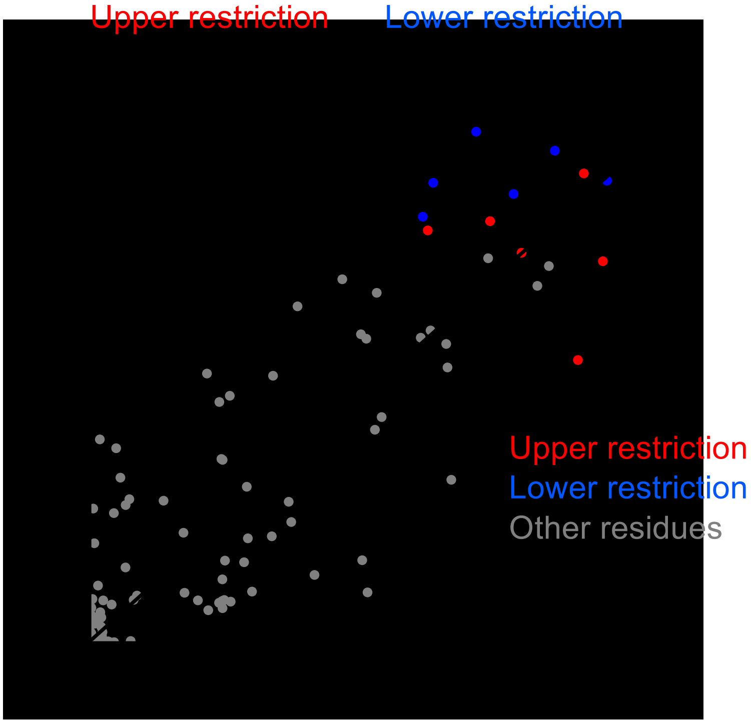
Upper restriction
Lower restriction
Upper restriction
Lower restriction
Other residues

**Supplementary Figure 7: Energetic restrictions within PTCH1 sterol transport tunnels.**

**A)** Conserved energetic peaks within the ECD (numbered 1-3 in Fig. 2D) of PTCH1-molA (yellow) and PTCH1-molB (light blue). The four residues with highest cholesterol interaction occupancy at each peak were identified from umbrella sampling windows using PyLipID[70] and a 0.6 nm cut-off. Residues are shown as spheres coloured by localisation within the upper (red) or lower (blue) restrictions in the PTCH1 ECD, surrounding putative ECD sterol transport tunnels (grey mesh). Residues corresponding to a particular peak are opaque and labelled, overlaid with those residues which comprise the remaining two peaks (transparent). **B)** Cholesterol interaction occupancy of equivalent residues in PTCH1-molA and PTCH1-molB across PMF-1a windows, indicating the same high occupancy residues contribute to formation of the upper (red) and lower (blue) restrictions between PTCH1 conformations. Cholesterol contacts were defined using a 0.6 nm cut-off.

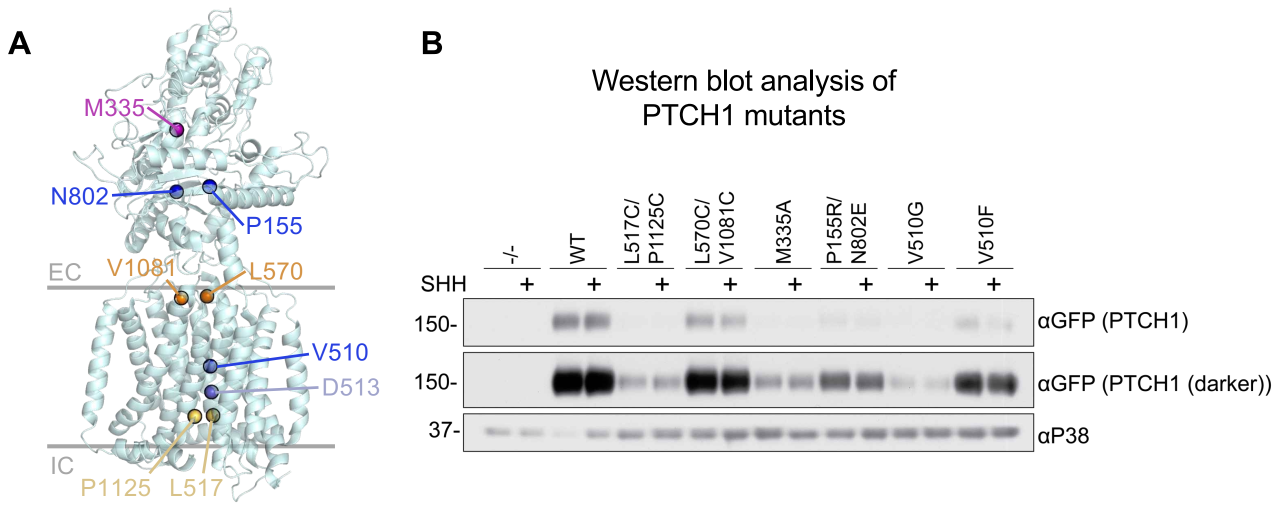


**Supplementary Figure 8: Expression levels of PTCH1 mutants.**
**A)** Location of PTCH1 mutants mapped onto the PTCH1 structure (PDB: 6DMY). **B)** Western blot analysis of PTCH1 WT and mutant variant expression levels in *Ptch1*$^{-/-}$ mouse embryonic fibroblasts in the presence or absence of SHH addition.

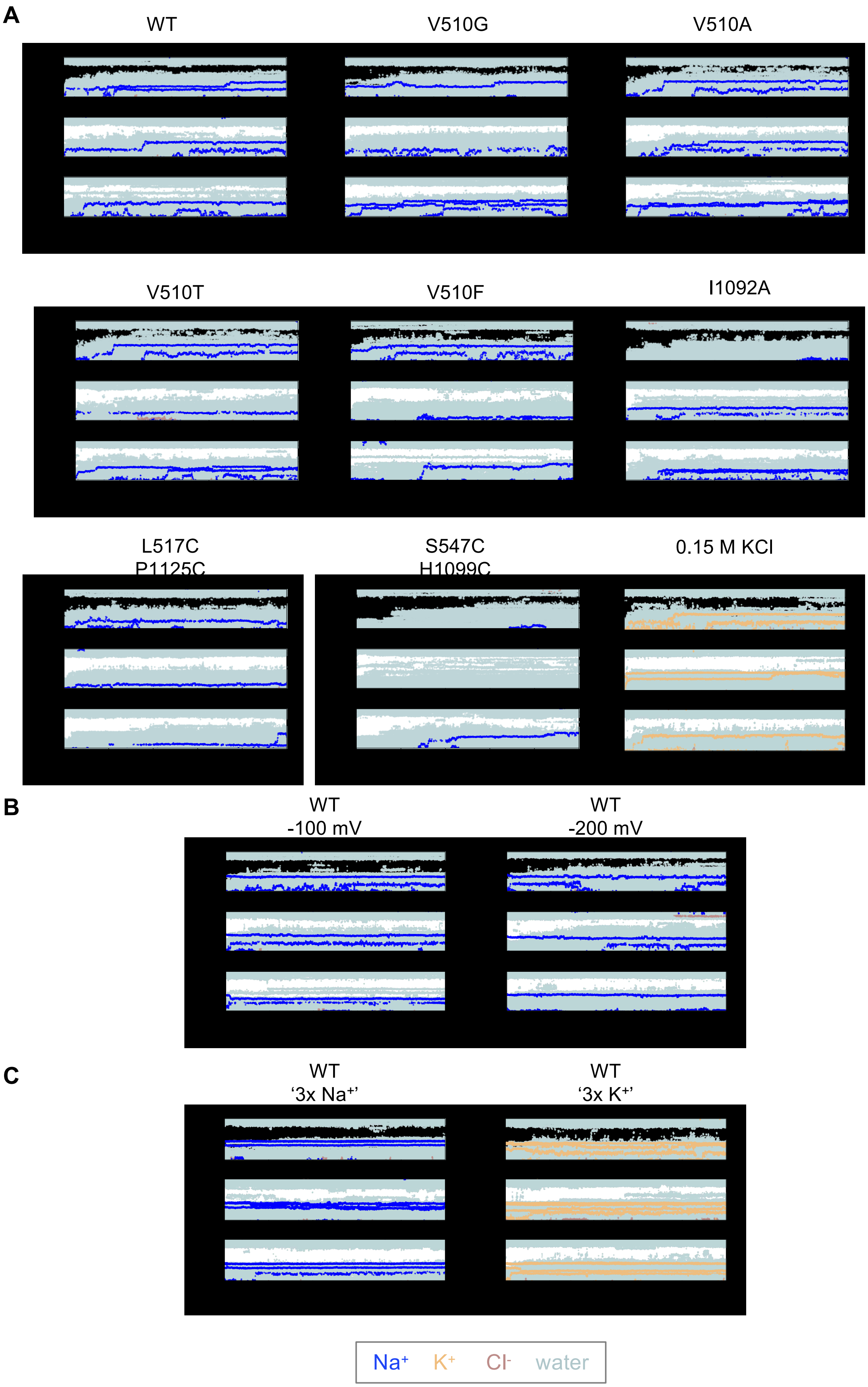

A
WT
V510G
V510A
V510T
V510F
I1092A
L517C
P1125C
S547C
H1099C
0.15 M KCl
B
WT
-100 mV
WT
-200 mV
C
WT
'3x Na+'
WT
'3x K+'
Na+
K+
Cl-
water

**Supplementary Figure 9: Water and ions within the TMD of PTCH1 mutants.**

**A)** The z coordinates of water oxygen atoms (light blue), $Na^+$ (blue), $K^+$ (yellow) and $Cl^-$ (salmon) ions localized within the PTCH1 TMD. A cylinder (length 4 nm, radius 1.3 nm) centred on the midpoint of V520 and I1092 Cα atoms was to identify water and ions within the PTCH1 TMD for WT and labelled PTCH1 mutants (V510G, V510A, V5105, V510F, I1092A, L517C/P1125C, S547C/H1099C). Simulations were initiated without $Na^+$/$K^+$ bound within the TMD and simulated for 3 x 50 ns or 3 x 100ns replicates. **B)** Water and ions within the WT PTCH1 TMD (defined as in **A**) accounting in the presence of a -100 mV or -200 mV membrane potential (see methods). **C)** As in **A-B** for WT PTCH1 simulations initiated with 3 x $Na^+$ or 3 x $K^+$ ions bound at equivalent positions to three $Na^+$ ions observed within a structure of DISP1 (PDB: 7RPH).

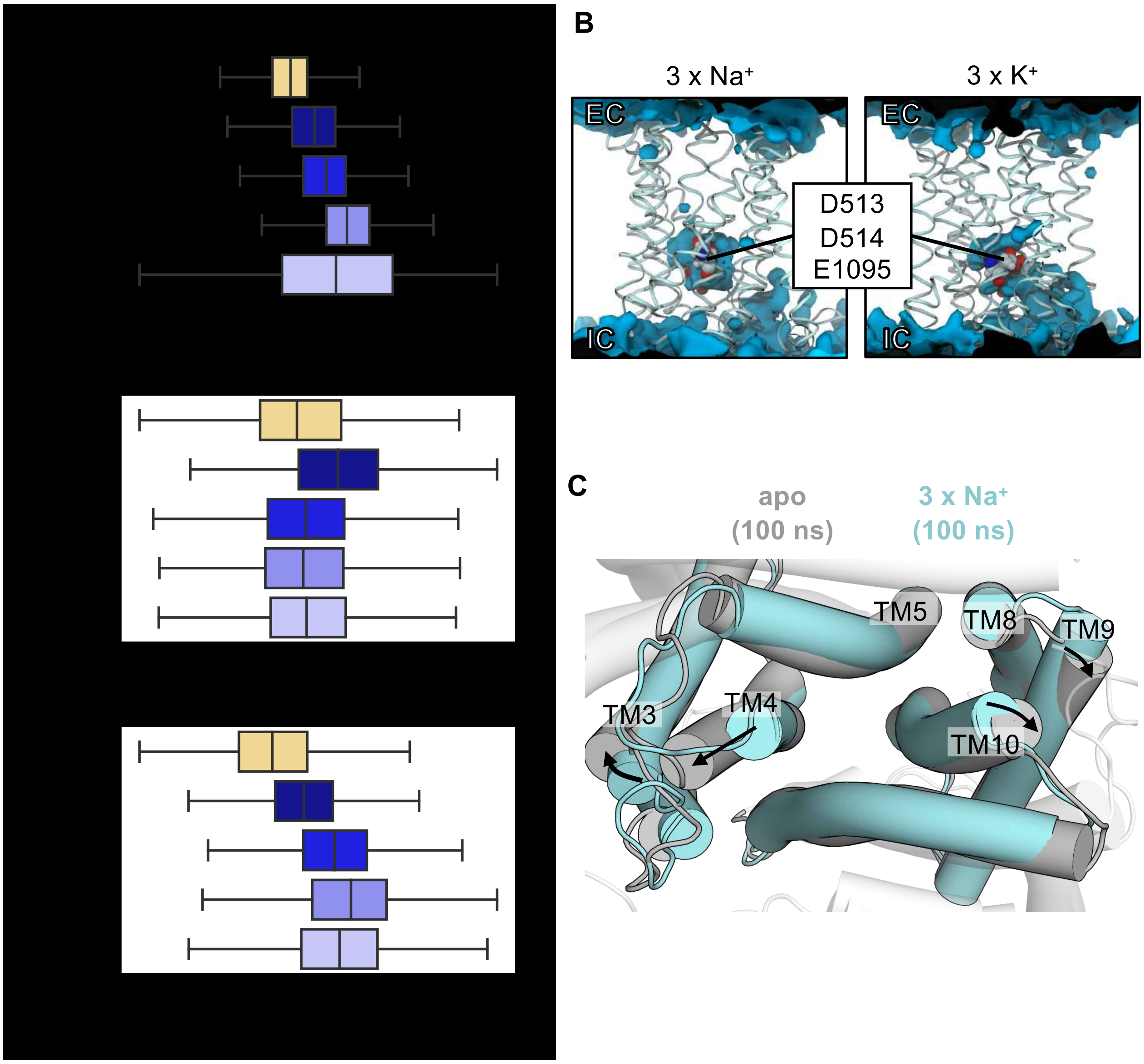


**Supplementary Figure 10: DISP1 extracellular TMD distances and PTCH1 TMD conformational changes.**

**A)** Minimum distances between the extracellular portions of transmembrane helices across 3 x 100 ns simulations of DISP1 in distinct ion coupled states. Distances were defined between the Cα atoms of A561-T1039 (TM4-TM10), V623-A985 (TM5-TM8) and Y542-V1022 (TM3-TM9). **B)** Time averaged water density (blue isosurface) across 100 ns simulations of PTCH1 (PDB: 6DMY) initiated with either 3 x $Na^+$ or 3 x $K^+$ ions bound within the TMD. Anionic triad residues are shown as spheres. **C)** Comparison of the intracellular PTCH1 TMD helices at the end of 100 ns atomistic simulations initiated in either an apo conformation or with 3x $Na^+$ ions bound within the TMD.
